# APPLES: Scalable Distance-based Phylogenetic Placement with or without Alignments

**DOI:** 10.1101/475566

**Authors:** Metin Balaban, Shahab Sarmashghi, Siavash Mirarab

## Abstract

Placing a new species on an existing phylogeny has increasing relevance to several applications. Placement can be used to update phylogenies in a scalable fashion and can help identify unknown query samples using (meta-)barcoding, skimming, or metagenomic data. Maximum likelihood (ML) methods of phylogenetic placement exist, but these methods are not scalable to reference trees with many thousands of leaves, limiting their ability to enjoy benefits of dense taxon sampling in modern reference libraries. They also rely on *assembled* sequences for the reference set and aligned sequences for the query. Thus, ML methods cannot analyze datasets where the reference consists of unassembled reads, a scenario relevant to emerging applications of genome-skimming for sample identification. We introduce APPLES, a distance-based method for phylogenetic placement. Compared to ML, APPLES is an order of magnitude faster and more memory efficient, and unlike ML, it is able to place on large backbone trees (tested for up to 200,000 leaves). We show that using dense references improves accuracy substantially so that APPLES on dense trees is more accurate than ML on sparser trees, where it can run. Finally, APPLES can accurately identify samples without assembled reference or aligned queries using kmer-based distances, a scenario that ML cannot handle. APPLES is available publically at github.com/balabanmetin/apples.

---

Phylogenetic placement is the problem of finding the optimal position for a new *query* species on an existing *backbone* (or, reference) tree. Placement, as opposed to a *de novo* reconstruction of the full phylogeny, has two advantages. In some applications (discussed below), placement is all that is needed, and in terms of accuracy, it is as good as, and even better than (Janssen *et al.*, 2018), *de novo* reconstruction. Moreover, placement can be more scalable than *de novo* reconstruction when dealing with very large trees.

Earlier research on placement was motivated by scalability. For example, placement is used in greedy algorithms that start with an empty tree and add sequences sequentially (e.g., Felsenstein, 1981; Desper and Gascuel, 2002). Each placement requires polynomial (often linear) time with respect to the size of the backbone, and thus, these greedy algorithms are scalable (often requiring quadratic time). Despite computational challenges (Warnow, 2017), there has been much progress in the *de novo* reconstruction of ultra-large trees (e.g., thousands to millions of sequences) using both maximum likelihood (ML) (e.g., Price *et al.*, 2010; Nguyen *et al.*, 2015) and the distance-based (e.g., Lefort *et al.*, 2015) approaches. However, these large-scale reconstructions require significant resources. As new sequences continually become available, placement can be used to update existing trees without repeating previous computations on full dataset.

More recently, placement has found a new application in sample identification: given one or more *query* sequences of unknown origins, detect the identity of the (set of) organism(s) that could have generated that sequence. These identifications can be made easily using sequence matching tools such as BLAST (Altschul *et al.*, 1990) when the query either exactly matches or is very close to a sequence in the reference library. However, when the sequence is novel (i.e., has lowered similarity to known sequences in the reference), this *closest* match approach is not sufficiently accurate (Koski and Golding, 2001), leading some researchers to adopt a phylogenetic approach (Sunagawa *et al.*, 2013; Nguyen *et al.*, 2014). Sample identification is essential to the study of mixed environmental samples, especially of the microbiome, both using 16S profiling (e.g., Gill *et al.*, 2006; Krause *et al.*, 2008) and metagenomics (e.g., von Mering *et al.*, 2007). It is also relevant to barcoding (Hebert *et al.*, 2003) and meta-barcoding (Clarke *et al.*, 2014; Bush *et al.*, 2017) and quantification of biodiversity (e.g., Findley *et al.*, 2013). Driven by applications to microbiome profiling, placement tools like pplacer (Matsen *et al.*, 2010) and EPA(-ng) (Berger *et al.*, 2011; Barbera *et al.*, 2019) have been developed. Researchers have also developed methods for aligning query sequence (e.g., Berger and Stamatakis, 2011; Mirarab *et al.*, 2012) and for downstream steps (e.g., Stark *et al.*, 2010; Matsen and Evans, 2013). These publications have made a strong case that for sample identification, placement is sufficient (i.e., *de novo* is not needed). Moreover, some studies (e.g., Janssen *et al.*, 2018) have shown that when dealing with fragmentary reads typically found in microbiome samples, placement can be *more* accurate than *de novo* construction and can lead to improved associations of microbiome with clinical information.

Existing phylogenetic placement methods have focused on the ML inference of the best placement – a successful approach, which nevertheless, suffers from two shortcomings. On the one hand, ML can only be applied when the reference species are *assembled* into full-length sequences (e.g., an entire gene) and are *aligned*; however, in new applications that we will describe, assembling (and hence aligning) the reference set is not possible. On the other hand, ML, while somewhat scalable, is still computationally demanding, especially in memory usage, and cannot place on backbone trees with many thousands of leaves. As the density of reference substantially impacts the accuracy and resolution of placement, this inability to use ultra-large trees as backbone also limits accuracy. This limitation has motivated alternative methods using local sensitive hashing (Brown and Truszkowski, 2013) and divide-and-conquer (Mirarab *et al.*, 2012).

Assembly-free and alignment-free sample identification using genome-skimming (Dodsworth, 2015) can also benefit from phylogenetic placement. A genome-skim is a shutgun sample of the genome sequenced at low coverage (e.g., 1X) – so low that assembling the nuclear genome is not possible (though, mitochondrial or plastid genomes can often be assembled). Genome-skimming promises to replace traditional marker-based barcoding of biological samples (Coissac *et al.*, 2016) but limiting analyses to organelle genome can limit resolution. Moreover, mapping reads to reference genomes is also possible only for species that have been assembled, which is a small fraction of the biodiversity on Earth. Sarmashghi *et al.* (2019) have recently shown that using shared *k*-mers, the distance between two unassembled genome-skims with low coverage can be accurately estimated. This approach, unlike assembling organelle genomes, uses data from the entire nuclear genome and hence promises to provide a higher resolution (e.g., at species or sub-species levels) while keeping the low sequencing cost. However, ML and other methods that require assembled sequences cannot analyze genome-skims, where both the reference and the query species are unassembled genome-wide bags of reads.

Distance-based approaches to phylogenetics are well-studied, but no existing tool can perform distance-based placement of a query sequence on a given backbone. The distance-based approach promises to solve both shortcomings of ML methods. Distance-based methods are computationally efficient and do not require assemblies. They only need distances (however computed). Thus, they can take as input assembly-free estimates of genomic distance estimated from low coverage genome-skims using Skmer (Sarmashghi *et al.*, 2019) or other alternatives (Haubold, 2014; Leimeister *et al.*, 2017; Yi and Jin, 2013; Benoit *et al.*, 2016; Fan *et al.*, 2015; Ondov *et al.*, 2016; Jain *et al.*, 2018). While alignment-based phylogenetics has been traditionally more accurate than alignment-free methods when both methods are possible, in these new scenarios, only alignment-free methods are applicable.

Here, we introduce a new method for distance-based phylogenetic placement called APPLES (Accurate Phylogenetic Placement using LEast Squares). APPLES uses dynamic programming to find the optimal distance-based placement of a sequence with running time and memory usage that scale linearly with the size of the backbone tree. We test APPLES in simulations and on real data, both for alignment-free and aligned scenarios.

## Materials and Methods

### Problem Statement

#### Notations

Let an unrooted tree *T* = (*V, E*) be a weighted connected acyclic undirected graph with leaves denoted by *ℒ* = {1 … *n*}. We let *T\** be the rooting of *T* on a leaf 1 obtained by directing all edges away from 1. For node *u* ∈ *V*, let *p*(*u*) denote its parent, *c*(*u*) denote its set of children, *sib*(*u*) denote its siblings, and *g*(*u*) denote the set of leaves at or below *u* (i.e., those that have *u* on their path to the root), all with respect to *T\**. Also let *l*(*u*) denote the length of the edge (*p*(*u*), *u*).

#### Distances

The tree *T* defines an *n* × *n* matrix where each entry *d*_*ij*_(*T*) corresponds to the path length between leaves *i* and *j*. We further generalize this definition so that *d*_*uv*_(*T \**) indicates the length of the undirected path between any two nodes of *T\** (when clear, we simply write *d*_*uv*_). Given some input data, we can compute a matrix of all pairwise sequence distances Δ, where the entry *δ*_*ij*_ indicates the dissimilarity between species *i* and When the sequence distance *δ*_*ij*_ is computed using (the correct) phylogenetic model, it will be a noisy but statistically consistent estimate of the tree distance *d*_*ij*_(*T*) (Felsenstein, 2003). Given these “phylogenetically corrected” distances (e.g 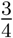 In 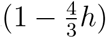 is the corrected hamming distance *h* under the Jukes and Cantor (1969) model), we can define optimization problems to recover the tree that best fits the distances. A natural choice is minimizing the (weighted) least square difference between tree and sequence distances:

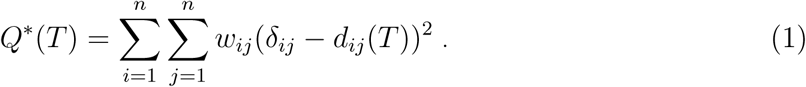

Here, weights (e.g., *w*_*ij*_) are used to reduce the impact of large distances (expected to have high variance). A general weighting schema can be defined as 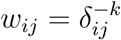 for a *constant* value *k* ∈ ℕ. Standard choices of *k* include *k* = 0 for the ordinary least squares (OLS) method of Cavalli-Sforza and Edwards 1967, *k* = 1 due to Beyer *et al.* 1974 (BE), and *k* = 2 due to Fitch and Margoliash 1967 (FM).

Finding arg min_*T*_ *Q\** (*T*) is NP-Complete (Day and Sankoff, 1987). However, decades of research has produced heuristics like neighbor-joining (Saitou and Nei, 1987), alternative formulations like (balanced) minimum evolution (Cavalli-Sforza and Edwards, 1967; Desper and Gascuel, 2002), and several effective tools for solving the problem heuristically (e.g., FastME by Lefort *et al.* 2015, DAMBE by Xia 2018, and Ninja by Wheeler 2009).

#### Phylogenetic placement

We let *P* (*u, x*_1_, *x*_2_) be the tree obtained by adding a query taxon *q* on an edge (*p*(*u*), *u*), creating three edges (*t, q*), (*p*(*u*), *t*), and (*t, u*), with lengths *x*_1_, *x*_2_, and *l*(*u*) − *x*_2_, respectively (Fig. 1). When clear, we simply write *P* and note that *P induces T* both in topology and branch length. We now define the problem.

**Fig. 1.**
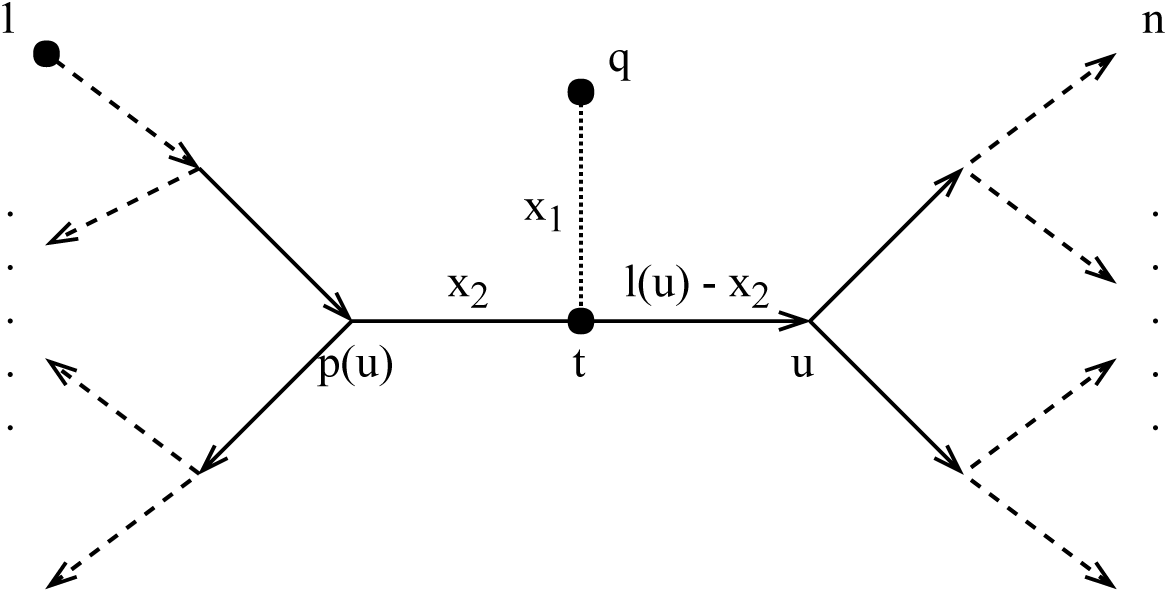
Any placement of *q* can be characterized as a tree *P* (*u, x*_1_, *x*_2_), shown here. The backbone tree *T \** is an arborescence on leaves *ℒ* = {1 …*n*}, rooted at leaf 1. Query taxon *q* is added on the edge between *u* and *p*(*u*), creating a node *t*. All placements on this edge are characterized by *x*_1_, the length of the pendant branch, and *x*_2_, the distance between *t* and *p*(*u*).

#### Least Squares Phylogenetic Placement (LSPP)

##### Input

A backbone tree *T* on *ℒ*, a query species *q*, and a vector Δ_*q\**_ with elements *δ*_*qi*_ giving sequence distances between *q* and every species *i* ∈ ***ℒ***;

##### Output

The placement tree *P* that adds *q* on *T* and minimizes

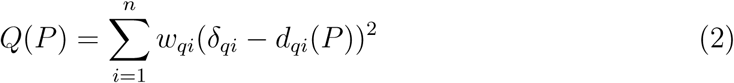

### Linear Time Solution for LSPP

The number of possible placements of *q* is 2*n* − 3. Therefore, LSPP can be solved by simply iterating over all the topologies, optimizing the score for that branch, and returning the placement with the minimum least square error. A naive algorithm can accomplish this in Θ(*n*^2^) running time by optimizing Eq. 2 for each of the 2*n* − 3 branches. However, using dynamic programming, the optimal solution can be found in linear time.

#### THEOREM 1

The LSPP problem can be solved with Θ(*n*) running time and memory.

The proof (given in Appendix A) follows easily from three lemmas that we next state. The algorithm starts with precomputing a fixed-size set of values for each nodes. For any node *u* and exponents *a* ∈ 𝕫 and *b* ∈ ℕ^+^, let 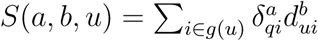 and for *b* = 0, let 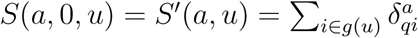. Note that *S′*(0, *u*) = |*g*(*u*)|. Similarly, for *u* ∈ *V* \ {1}, let 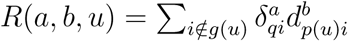 for *b >* 0 and let 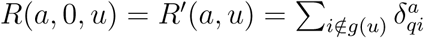.

#### LEMMA 2

The set of all *S*(*a, b, u*) and *R*(*a, b, u*) values can be precomputed in Θ(*n*) time with two tree traversals using the dynamic programming given by:

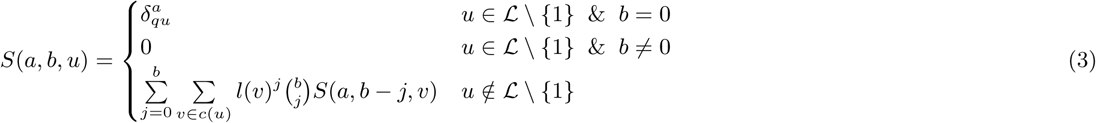

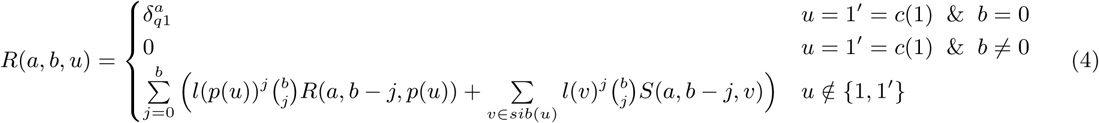

#### LEMMA 3

Equation 2 can be rearranged (see Eq. S2 in Appendix A) such that computing *Q*(*P*) for a given *P* = *P* (*u, x*_1_, *x*_2_) requires a constant time computation using *S*(*a, b, u*) and *R*(*a, b, u*) values for −*k* ≤ *a* ≥ 2 − *k* and 0 ≤ *b* ≥ 2.

Thus, after a linear time precomputation, we can compute the error for any given placement in constant time. It remains to show that for each node, the optimal placement on the branch above it (e.g., *x*_1_ and *x*_2_) can be computed in constant time.

#### LEMMA 4

For a fixed node *u* ∈ *V* \ {1}, if 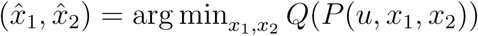, then

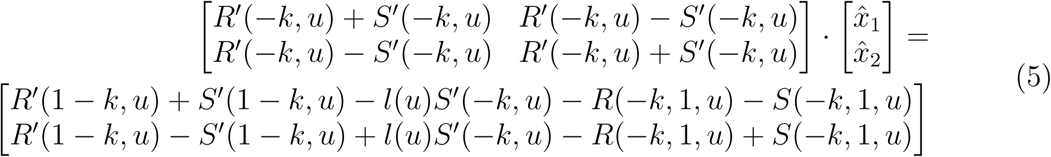

and hence 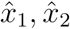 can be computed in constant time.

#### Non-negative branch lengths

The solution to Equation 5 does not necessarily conform to constraints 0 ≤ *x*_1_ and 0 ≤ *x*_2_ ≤ *l*(*u*). However, the following lemma (proof in Appendix A) allows us to easily impose the constraints by choosing optimal boundary points when unrestricted solutions fall outside boundaries.

##### LEMMA 5

With respect to variables *x*_1_ and *x*_2_, *Q*(*P* (*u, x*_1_, *x*_2_)) is a convex function.

#### Minimum evolution

An alternative to directly using MLSE (Eq. 1) is the minimum evolution (ME) principle (Cavalli-Sforza and Edwards, 1967; Rzhetsky and Nei, 1992). Our algorithm can also optimize the ME criterion: after computing *x*_1_ and *x*_2_ by optimizing MLSE for each node *u*, we choose the placement with the minimum total branch length. This is equivalent to using arg min_*u*_ *x*_1_, since the value of *x*_2_ does not contribute to total branch length. Other solutions for ME placement exist (Desper and Gascuel, 2002), a topic we return to in the Discussion section.

#### Hybrid

We have observed cases where ME is correct more often than MLSE, but when it is wrong, unlike MLSE, it has a relatively high error. This observation led us to design a hybrid approach. After computing *x*_1_ and *x*_2_ for all branches, we first select the top *log*_2_(*n*) edges with minimum *Q*(*P* (*u, x*_1_, *x*_2_)) values (this requires Θ(*n* log log *n*) time). Among this set of edges, we place the query on the edge satisfying the ME criteria.

### Datasets

We benchmark accuracy and scalability of APPLES in two settings: sample identification using assembly-free genome-skims on real biological data and placement using aligned sequences on simulated data.

### Real genome-skim datasets for the assembly-free scenario

#### Columbicola genome-skims

We use a set of 61 genome-skims by Boyd *et al.* (2017), including 45 known lice species (some represented multiple times) and 7 undescribed species. We generate lower coverage skims of 0.1Gb or 0.5Gb by randomly subsampling the reads from the sequence read archives (SRA) provided by the original publication (NCBI BioProject PRJNA296666). We use BBTools (Bushnell, 2014) to filter subsampled reads for adapters and contaminants and remove duplicated reads. Since this dataset is not assembled, the coverage of the genome-skims is unknown; Skmer estimates the coverage to be between 0.2X and 1X for 0.1Gb samples (and 5 times that coverage with 0.5Gb).

#### Anopheles and Drosophila datasets

We also use two insect datasets used by Sarmashghi *et al.* (2019): a dataset of 22 *Anopheles* and a dataset of 21 *Drosophila* genomes (Table S1), both obtained from InsectBase (Yin *et al.*, 2016). For both datasets, genome-skims with 0.1Gb and 0.5Gb sequence were generated from the assemblies using the short-read simulator tool ART, with the read length *l* = 100 and default error profile. Since species have different genome sizes, with 0.1Gb data, our subsampled genome-skims range in coverage from 0.35X to 1X for Anopheles and from 0.4X to 0.8X for Drosophila.

More recently, Miller *et al.* (2018) sequenced several Drosophila genomes, including 12 species shared with the InsectBase dataset. Sarmashghi *et al.* (2019) subsampled the SRAs from this second project to 0.1Gb or 0.5Gb and, after filtering contaminants, obtained artificial genome-skims. We can use these genome-skims as query and the genome-skims from the InsectBase dataset as the backbone. Since the reference and query come from two projects, the query genome-skim can have a non-zero distance to the same species in the reference set, providing a realistic test of sample identification applications.

### Simulated datasets for the aligned sequence scenario

#### GTR

We use a 101-taxon dataset available from Mirarab and Warnow 2015. Sequences were simulated under the General Time Reversible (GTR) plus the Γ model of site rate heterogeneity using INDELible (Fletcher and Yang, 2009) on gene trees that were simulated using SimPhy (Mallo *et al.*, 2016) under the coalescent model evolving on species trees generated under the Yule model. Note that the same model is used for inference under ML placement methods (i.e., no model misspecification). We took all 20 replicates of this dataset with mutation rates between 5 × 10^−8^ and 2 × 10^−7^, and for each replicate, randomly selected five estimated gene trees among those with ≤20% RF distance between estimated and true gene tree. Thus, we have a total of 100 backbone trees. This dataset is the simplest test case where model violation or miss-alignment is not a concern.

#### RNASim

Guo *et al.* 2009 designed a complex model of RNA evolution that does not make usual i.i.d assumptions of sequence evolution. Instead, it uses models of energy of the secondary structure to simulate RNA evolution by a mutation-selection population genetics model. This model is based on an inhomogeneous stochastic process without a global substitution matrix. The model complexity of RNASim allows us to test both ML and APPLES under a substantially misspecified model. An RNASim dataset of one-million sequences (with E.coli SSU rRNA used as the root), is available from Mirarab *et al.* 2015. We created several subsets of the full RNASim dataset.

i. Heterogeneous: We first randomly subsampled the full dataset to create 10 datasets of size 10, 000. Then, we chose the largest clade of size at most 250 from each replicate; this gives us 10 backbone trees of mean size 249.
ii. Varied diameter: To evaluate the impact of the evolutionary diameter (i.e., the highest distance between any two leaves in the backbone), we also created datasets with low, medium, and high diameters. We sampled the largest five clades of size at most 250 from each of the 10 replicates used for the heterogeneous dataset. Among these 50 clades, we picked the bottom, middle, and top five clades in diameter, which had diameter in [0.3, 0.4] (mean: 0.36), [0.5, 0.52] (mean: 0.51), and [0.65, 1.07] (mean: 0.82), respectively.
iii. Varied size (RNASim-VS): We randomly subsampled the tree of size one-million to create 5 replicates of datasets of size (*n*): 500, 1000, 5000, 10000, 50000, and 100000, and 1 replicate (due to size) of size 200000. For replicates that contain at least 5000 species, we removed sites that contain gaps in 95% or more of the sequences in the alignment.
iv. Query scalability (RNASim-QS): We first randomly subsampled the full one-million sequence RNASim dataset to create a dataset of size 500. Then for *k* =1 to 49,152 queries (choosing all *k* = 3 × 2^*i*^, 0 ≤ *i* ≤ 14) we created 5 replicates of *k* query sequences, again randomly subsampling from the full alignment with one-million sequences.
v. Alignment Error (RNASim-AE): Mirarab *et al.* (2015) used PASTA to estimate alignments on subsets of the RNASim dataset with up to 200,000 sequences. We use their reported alignment with 200,000 or 10,000 sequences (taking only replicate 1 in this case).

### Methods

#### Alternative methods

For aligned data, we compare APPLES to two ML methods: pplacer (Matsen *et al.*, 2010) and EPA-ng (Barbera *et al.*, 2019). Matsen *et al.* (2010) found pplacer to be substantially faster than EPA (Berger and Stamatakis, 2011) while their accuracy was similar. EPA-ng improves the scalability of EPA; thus, we compare to EPA-ng in analyses that concerned scalability (e.g., RNASim-VS). We run pplacer and EPA-ng in their default mode using GTR+Γ model and use their best hit (ML placement). We also compare with a simple method referred to as CLOSEST that places the query as the sister to the species with the minimum distance to it. CLOSEST is meant to emulate the use of BLAST (if it could be used). For the assembly-free setting, existing phylogenetic placement methods cannot be used, and we compared only against CLOSEST.

#### APPLES Software

We implemented APPLES using python, internally using Treeswift (Moshiri, 2018) for phylogenetic operations. APPLES computes distances internally (for certain models described below) using vectorized numpy (Oliphant, 2006) operations but can also use input distance matrices (e.g. generated using FastME). It generates the output in the standard jplace format (Matsen *et al.*, 2012).

#### Distance calculation and models

In computing pairwise distances, positions that have a gap in at least one of the two sequences are always ignored. In our analyses, we exclusively use models for nucleotide sequences. We compute phylogenetic distances under the parameter-free JC69 model, the six-parameter Tamura and Nei 1993 (TN93) model, and the 12-parameter general Markov model (Lockhart *et al.*, 1994). We compute distances independently for all pairs, and not simultaneously as suggested by Tamura *et al.* (2004). We also use the Gamma model of sites rate heterogeneity for JC69 and TN93 using the standard approach (Waddell and Steel, 1997). Pairing Gamma with GTR is theoretically possible in the absence of noise; however, the method can run into problems on real data (Waddell and Steel, 1997). Thus, we do not include a GTR model directly. Instead, we use the log-det approach that can handle the most general (12-parameter) Markov model (Lockhart *et al.*, 1994); however, log-det cannot account for rate across sites heterogeneity (Waddell and Steel, 1997). The *α* parameter of the Gamma model cannot be computed from pairwise sequence comparisons (Steel, 2009); instead, we use the *α* computed from the backbone tree. We used the *α* parameter computed by RAxML (Stamatakis, 2014) run on the backbone alignment and given the backbone tree. For analyses with models other than JC69, we use FastME to compute distances (see Supplementary material for details) while for JC69, distances are computed by APPLES.

In analyses on assembly-free datasets, we first compute genomic distances using Skmer (Sarmashghi *et al.*, 2019). We then correct these distances using the JC69 model, without the Gamma model of rate variation.

#### APPLES parameters

We have chosen default parameter settings for APPLES and refer to this version as APPLES*\**. By default, we use FM weighting, the MLSE selection criterion, enforcement of non-negative branch lengths, and JC69 distances. When not specified otherwise, these default parameters are used.

#### Backbone alignments and trees

For genome-skimming experiments, we estimated the backbone tree using FastME from the JC69 distance matrix computed from genome-skims using Skmer. By design, no alignments are needed.

For simulated datasets, we present results both based on true backbone alignments (for all datasets) and estimated alignments (for large datasets). The true alignments are known from the simulations. To test the accuracy in the presence of alignment error, we use the available PASTA backbone alignment for RNASim-AE dataset. The alignments have considerable error as measured by FastSP (Mirarab and Warnow, 2011): 11.5% and 12.7% SPFN, 10.9% and 11.7% SPFP, and only 165 and 848 fully correctly aligned sites (2.2% and 6.4%), respectively for 10,000 and 200,000 sequences. As before, here, we remove sites with more than 95% gaps in the estimated alignments.

For true alignments, we ran RAxML (Stamatakis, 2014) using GTRGAMMA model for all datasets to compute the topology of the backbone tree except for RNASim-AE and RNASim-VS dataset, where due to the size, we used FastTree-2 (Price *et al.*, 2010). When estimated alignment is used (RNASim-AE dataset), we used the co-estimated tree output by PASTA, which is itself computed using FastTree-2. We always re-estimated branch lengths and model parameters on that fixed topology using RAxML (switching to GTRCAT for *n* ≥ 10, 000) before running ML methods. For APPLES, we re-estimated branch lengths using FastTree-2 under the JC69 model to match the model used for estimating distances.

#### Alignment of queries

For analyses with true alignment as the backbone, we also use the true alignment of the queries to the backbone. In analyses with estimated backbone alignments, we align the query sequences to the estimated backbone alignment using SEPP (Mirarab *et al.*, 2012), which is a divide-and-conquer method that internally uses HMMER (Eddy, 1998, 2009), with alignment subset size set to 10% of the full set (default setting). We use the resulting extended alignment, after masking unaligned sites, to run both APPLES and EPA-ng, in both cases, placing on the full backbone tree. We also report results using default SEPP (which runs pplacer internally); however, here, due to limitations of pplacer, we use the default setting of SEPP, which set the placement subset size to 10% of the full set.

### Evaluation Procedure

To evaluate the accuracy, we use a leave-one-out strategy. We remove each leaf *i* from the backbone tree *T* and place it back on this *T* \ *i* tree to obtain the placement tree *P*. On the RNAsim-VS dataset, due to its large size, we only removed and added back 200 randomly chosen leaves per replicate. On the RNAsim-AE dataset, we remove 200 queries from the backbone at the same time (leave-many-out). Finally, for RNAsim-QS, we place *k* queries *in one run*, allowing the methods to benefit from optimizations designed for multiple queries, but note that queries are not selected from the backbone tree but are instead selected from the full dataset. In all cases, placement of queries is with respect to the backbone and not other queries.

#### Delta error

We measure the accuracy of a placement tree *P* of a single query *q* on a backbone tree *T* on leafset *ℒ* with respect to the true tree *T\** on *ℒ∪* {*q*} using delta error:

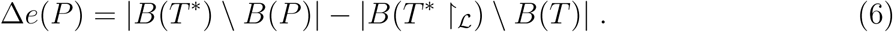

where *B*(.) is the set of bipartitions of a tree and *T\** ↾_*ℒ*_ is the true tree restricted to *ℒ*. Note that Δ*e*(*P*) ≥ 0 because adding *q* cannot decrease the number of missing branches in *T*. We report delta error averaged over all queries (denoted as Δ*e*). In leave-one-out experiments, placing *q* to the same location as the backbone before leaving it out can still have a non-zero delta error because the backbone tree is not the true tree. We refer to the placement of a leaf into its position in the backbone tree as the *de novo* placement. In leave-many-out experiments, we measure delta error of each query separately (not the delta error of the combination of all queries). On biological data, where the true tree is unknown, we use a reference tree (Fig. S1). For Drosophila and Anopheles, we use the tree available from the Open Tree Of Life (Hinchliff *et al.*, 2015) as the reference. For Columbicola, we use the ML concatenation tree published by Boyd *et al.* (2017) as the reference.

## Results

### Assembly-free Placement of Genome-skims

On our three biological genome-skim datasets, APPLES*\** successfully places the queries on the optimal position in most cases (97%, 95%, and 71% for Columbicola, Anopheles, and Drosophila, respectively) and is never off from the optimal position by more than one branch. Other versions of APPLES are less accurate than APPLES*\**; e.g., APPLES with ME can have up to five wrong branches (Table 1). On genome-skims, where assembly and alignment are not possible, existing placement tools cannot be used, and the only alternative is the CLOSEST method (emulating BLAST if assembly was possible). CLOSEST finds the optimal placement only in 54% and 57% of times for Columbicola and Drosophila; moreover, it can be off from the best placement by up to seven branches for the Columbicola dataset. On the Anopheles dataset, where the reference tree is unresolved (Fig. S1), all methods perform similarly.

**Table 1.** Assembly-free placement of genome-skims. We show the percentage of placements into optimal position (those that do not increase Δ*e*), average delta error (Δ*e*), and maximum delta error (*e*_*max*_) for APPLES, assignment to the CLOSEST species, and the placement to the position in the backbone (DE-NOVO) over the 61 (a), 22 (b), and 21 (c) placements. Results are shown for genome skims with 0.1Gbp of reads. Delta error is the increase in the missing branches between the reference tree and the backbone tree after placing each query.

|  | (a) *Columbicola* |  |  | (b) *Anopheles* |  |  | (c) *Drosophila* |  |  |
| --- | --- | --- | --- | --- | --- | --- | --- | --- | --- |
| | % | $\Delta e$ | $e_{max}$ | % | $\Delta e$ | $e_{max}$ | % | $\Delta e$ | $e_{max}$ |
| **APPLES\*** | 97 | 0.03 | 1 | 95 | 0.05 | 1 | 71 | 0.29 | 1 |
| **APPLES-ME** | 84 | 0.28 | 5 | 95 | 0.05 | 1 | 67 | 0.42 | 2 |
| **APPLES-HYBRID** | 87 | 0.16 | 2 | 95 | 0.05 | 1 | 67 | 0.33 | 1 |
| **CLOSEST** | 54 | 1.15 | 7 | 91 | 0.09 | 1 | 57 | 0.62 | 3 |
| **DE-NOVO** | 98 | 0.02 | 1 | 95 | 0.05 | 1 | 71 | 0.29 | 1 |

APPLES*\** is less accurate on the Drosophila dataset than other datasets. However, here, simply placing each query on its position in the backbone tree would lead to identical results (Table 1). Thus, placements by APPLES*\** are as good as the *de novo* construction, meaning that errors of APPLES*\** are entirely due to the differences between our backbone tree and the reference tree. Moreover, these errors are not due to low coverage; increasing the genome-skim size 5x (to 0.5Gb) does not decrease error (Table S4).

On Drosophila dataset, we next tested a more realistic sample identification scenario using the 12 genome-skims from the separate study (and thus, non-zero distance to the corresponding species in the backbone tree). As desired, APPLES*\** places all of 12 queries from the second study as sister to the corresponding species in the reference dataset.

### Alignment-based Placement

We first compare the accuracy and scalability of APPLES*\** to ML methods and then compare various settings of APPLES. For ML, we use pplacer (shown everywhere) and EPA-ng (shown only when we study scalability and work on large backbones).

### Comparison to Maximum Likelihood (ML) without Alignment Error

#### GTR dataset

On this dataset, where it faces no model misspecification, pplacer has high accuracy. It finds the best placement in 84% of cases and is off by one edge in 15% (Fig. 2a); its mean delta error (Δ*e*) is only 0.17 edges. APPLES*\** is also accurate, finding the best placement in 78% of cases and resulting in the mean Δ*e* =0.28 edges. Thus, even though pplacer uses ML and faces no model misspecification and APPLES*\** uses distances based on a simpler model, the accuracy of the two methods is within 0.1 edges on average. In contrast, CLOSEST has poor accuracy and is correct only 50% of the times, with the mean Δ*e* of 1.0 edge.

**Fig. 2.**
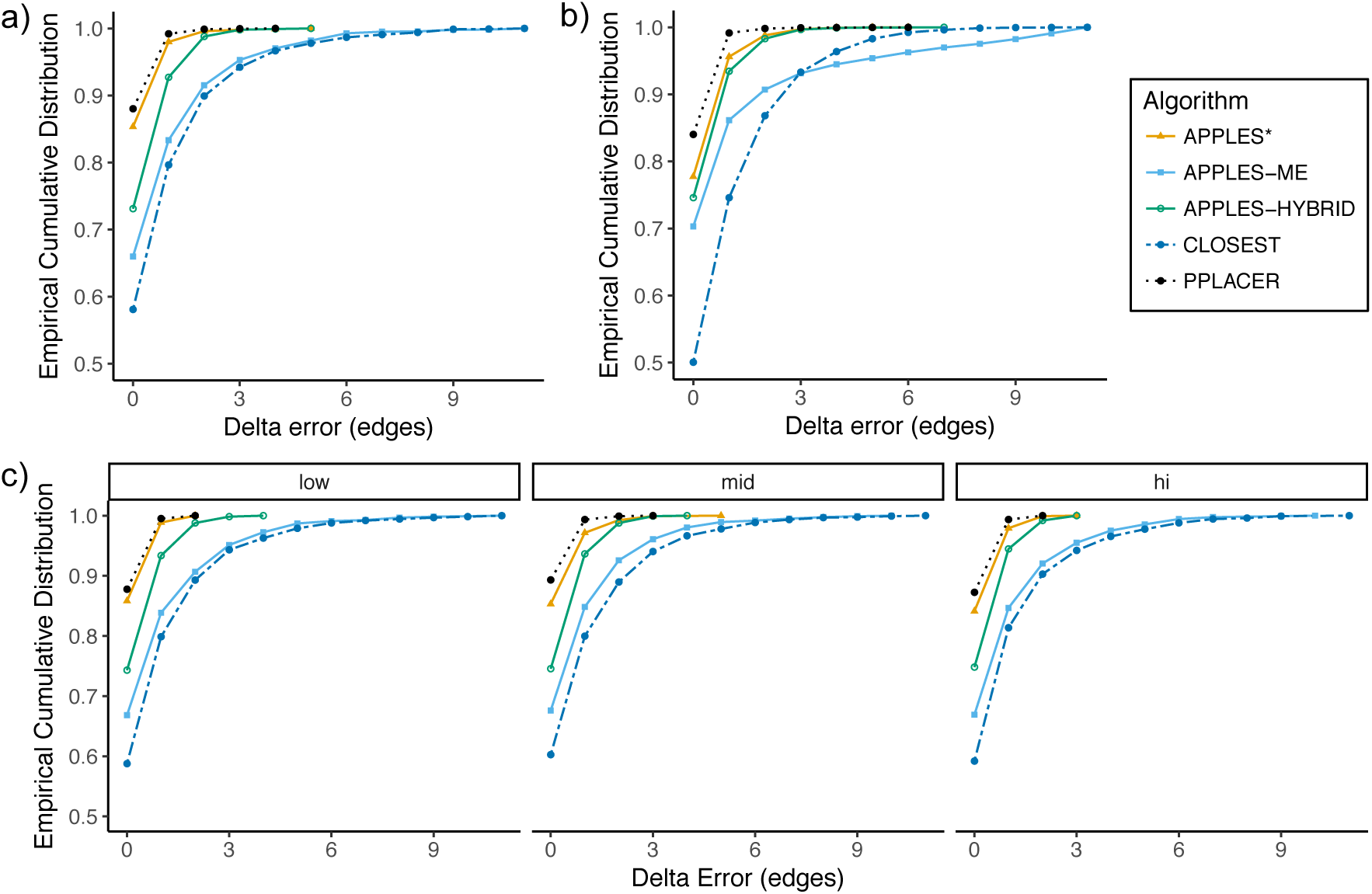
Accuracy on simulated data. We show empirical cumulative distribution of the delta error, defined as the increase in the number of missing branches in the estimated tree compared to the true tree. We compare pplacer (dotted), CLOSEST match (dashed), and APPLES with FM weighting and JC69 distances and MLSE (APPLES$), ME, or Hybrid optimization. (a) GTR dataset. (b) RNASim-Heterogeneous. (c) RNASim-varied diameter, shown in boxes: low, medium (mid), or high. Distributions are over 10, 000 (a), 2450 (b), and 3675 (c) points.

#### Model misspecification

On the small RNASim data with subsampled clades of ≈ 250 species), both APPLES*\** and pplacer face model misspecification. Here, the accuracy of APPLES*\** is very close to ML using pplacer. On the heterogeneous subset (Fig. 2b and Table 2), pplacer and APPLES*\** find the best placement in 88% and 85% of cases and have a mean delta error of 0.13 and 0.17 edges, respectively. Both methods are much more accurate than CLOSEST, which has a delta error of 0.87 edges on average.

**Table 2.**
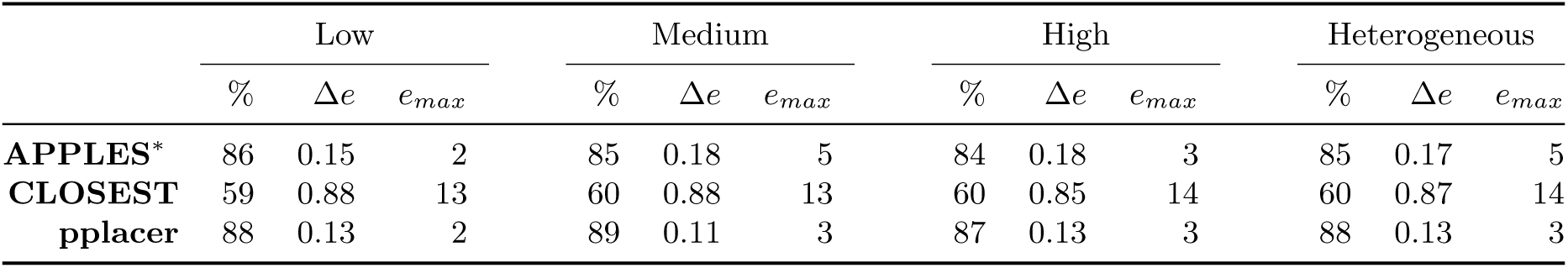
The delta error for APPLES*\**, CLOSEST match, and pplacer on the RNASim-varied diameter dataset (low, medium, or high) and the RNA-heterogeneous dataset. Measurements are shown over 1250 placements for each diameter size category, corresponding to 5 backbone trees and 250 placements per replicate.

|  | Low |  |  | Medium |  |  | High |  |  | Heterogeneous |  |  |
| --- | --- | --- | --- | --- | --- | --- | --- | --- | --- | --- | --- | --- |
| | % | $\Delta e$ | $e_{max}$ | % | $\Delta e$ | $e_{max}$ | % | $\Delta e$ | $e_{max}$ | % | $\Delta e$ | $e_{max}$ |
| **APPLES\*** | 86 | 0.15 | 2 | 85 | 0.18 | 5 | 84 | 0.18 | 3 | 85 | 0.17 | 5 |
| **CLOSEST** | 59 | 0.88 | 13 | 60 | 0.88 | 13 | 60 | 0.85 | 14 | 60 | 0.87 | 14 |
| **pplacer** | 88 | 0.13 | 2 | 89 | 0.11 | 3 | 87 | 0.13 | 3 | 88 | 0.13 | 3 |

#### Impact of diameter

When we control the tree diameter, APPLES*\** and pplacer remain very close in accuracy (Fig. 2c). The changes in error are small and not monotonic as the diameters change (Table 2). The accuracies of the two methods at low and high diameters are similar. The two methods are most divergent in the medium diameter case, where pplacer has its lowest error (Δ*e* =0.11) and APPLES*\** has its highest error (Δ*e* =0.18).

To summarize results on small RNASim dataset with model misspecification, although APPLES*\** uses a parameter-free model, its accuracy is extremely close to ML using pplacer with the GTR+Γ model.

#### Impact of taxon sampling

The real advantage of APPLES*\** over pplacer becomes clear for placing on larger backbone trees (Fig. 3 and Table 3). For backbone sizes of 500 and 1000, pplacer continues to be slightly more accurate than APPLES*\** (mean Δ*e* of pplacer is better than APPLES*\** by 0.09 and 0.23 edges, respectively). However, with backbones of 5000 leaves, pplacer fails to run on 449/1000 cases, producing infinity likelihood (perhaps due to numerical issues) and has 41 times higher error than APPLES*\** on the rest (Fig. S2). Since pplacer could not scale to 5,000 leaves, we also test the recent method, EPA-ng (Barbera *et al.*, 2019). Given the 64GB of memory available on our machine, EPA-ng is able to run on datasets with up to 10, 000 leaves. EPA-ng finds the correct placement less often than pplacer but is close to APPLES*\** (Fig. 3a). However, when it has error, it tends to have a somewhat lower distance to the correct placement, making its mean error slightly better than APPLES*\** (Fig. 3b and Table 3).

**Table 3.** Percentage of correct placements (shown as %) and the delta error (Δ*e*) on the RNASim datasets with various backbone size (*n*). % and Δ*e* is over 1000 placements (except *n* = 200, 000, which is over 200 placements). Running pplacer and EPA-ng was not possible (n.p) for trees with at least 10, 000 leaves and failed in some cases (number of fails shown) for 5, 000 leaves.

|  | n = 500 |  | n = 1,000 |  | n = 5,000 |  | n = 10000 |  | n = 100,000 |  | n = 200,000 |  |
| --- | --- | --- | --- | --- | --- | --- | --- | --- | --- | --- | --- | --- |
| | % | $\Delta e$ | % | $\Delta e$ | % | $\Delta e$ | % | $\Delta e$ | % | $\Delta e$ | % | $\Delta e$ |
| **APPLES\*** | 75 | 0.32 | 71 | 0.43 | 77 | 0.37 | 79 | 0.33 | 84 | 0.25 | 87 | 0.25 |
| **CLOSEST** | 52 | 1.16 | 53 | 1.18 | 54 | 1.15 | 59 | 0.90 | 61 | 0.69 | 63 | 0.70 |
| **EPA-ng** | 73 | 0.33 | 73 | 0.31 | 78 | 0.24 | 79 | 0.22 | n.p | n.p | n.p | n.p |
| **pplacer** | 80 | 0.23 | 81 | 0.20 | fail (800) | n.p | n.p | n.p | n.p | n.p | n.p | n.p |

**Fig. 3.**
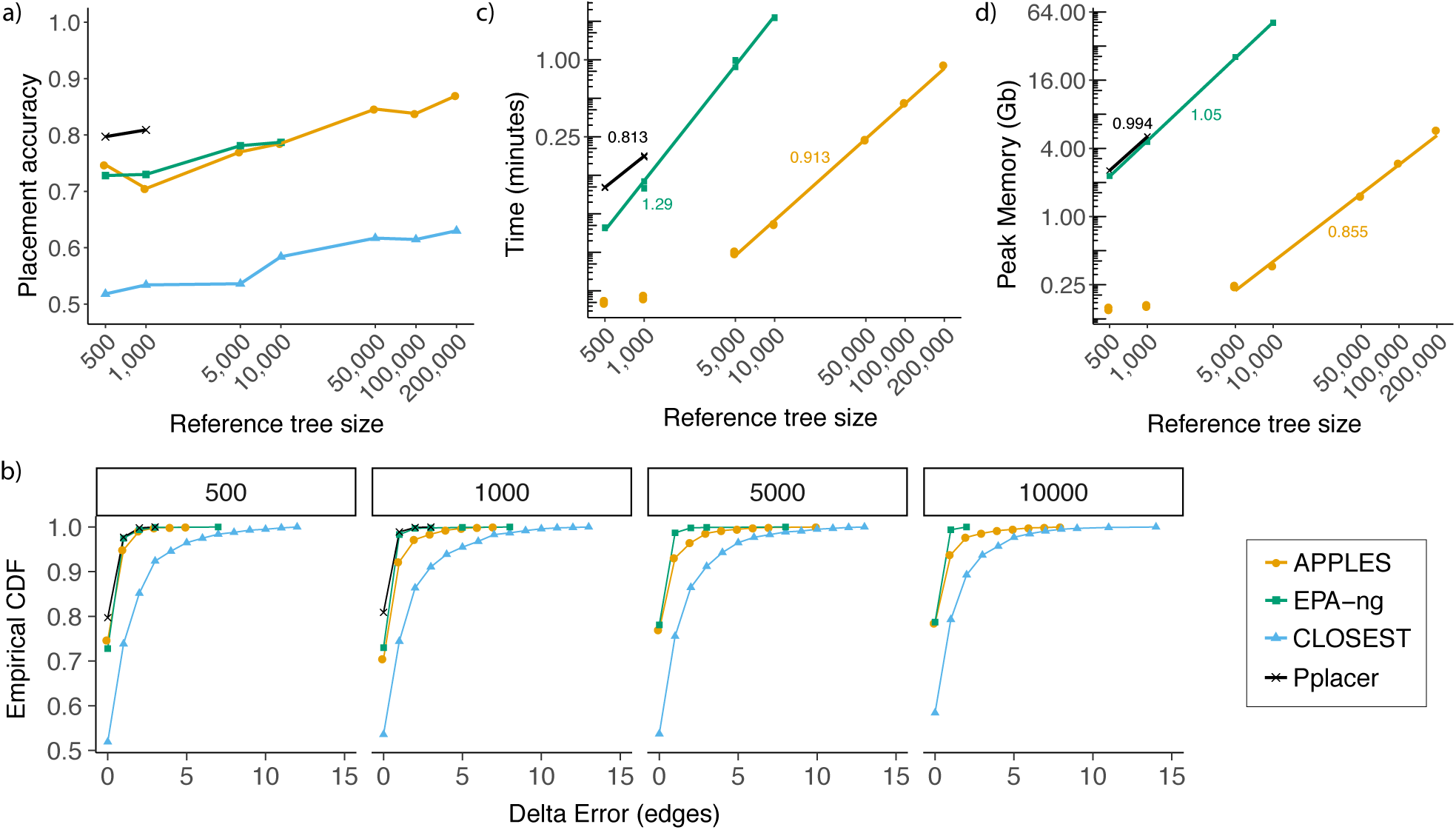
Results on RNASim-VS. (a) Placement accuracy with taxon sampling ranging from 500 to 200,000. (b) The empirical cumulative distribution of the delta error, shown for 500 ≤ *n* ≤ 10000 where EPA-ng can run. (c,d) Running time and peak memory usage of placement methods for a single placement. Lines are fitted in the log-log scale and their slope (indicated on the figure) empirically estimates the polynomial degree of the asymptotic growth (*t* = *n*^*a*^ ⇒ log *t* = *a* log *n*). APPLES lines are fitted to ≥ 5, 000 points because the first two values are small and irrelevant to asymptotic behavior. All calculations are on 8-core, 2.6GHz Intel Xeon CPUs (Sandy Bridge) with 64GB of memory, with each query placed independently and given 1 CPU core and the entire memory.

For backbones trees with more than 10, 000 leaves, pplacer and EPA-ng are not able to run given computational resources at hand, and CLOSEST is not very accurate (finding the best placement in only 59% of cases). However, APPLES*\** continues to be accurate for all backbone sizes. As the backbone size increases, the taxon sampling of the tree is improving (recall that these trees are all random subsets of the same tree). With denser backbone trees, APPLES*\** has increased accuracy despite placing on larger trees (Fig. 3a, Table 3). For example, using a backbone tree of 200, 000 leaves, APPLES*\** is able to find the best placement of query sequences in 87% of cases, which is better than the accuracy of either APPLES*\** or ML tools on any backbone size. Thus, an increased taxon sampling helps accuracy, but ML tools are limited in the size of the tree they can handle given relatively powerful machines (e.g., 64GB of memory).

#### Running time and memory

As the backbone size increases, the running times of pplacer and APPLES grow close to linearly with the size of the backbone tree, *n*, whereas running time of EPA-ng seems to grow with *n*^1.3^ (Fig. 3c). APPLES is on average 13 times faster than pplacer and 7.5 times faster than EPA-ng on backbone trees with 1000 leaves, and is 41 times faster than EPA-ng with 10,000-taxon backbones.

The memory consumption of all methods increases close to linearly with the *n*, but APPLES requires dramatically less memory (Fig. 3d). For example, for placing on a backbone with 10,000 leaves, EPA-ng requires 51GB of memory, whereas APPLES requires only 0.4GB. APPLES easily scales to a backbone of 200, 000 sequences, running in only 1 minute and using 6GB of memory per query (including all precomputations in the dynamic programming). These numbers also include the time and memory needed to compute the distance between the query sequence and all the backbone sequences.

We next test the scalability of APPLES*\** and EPA-ng with respect to the number of queries, *k*, in one run given 28 CPU cores on RNASim-QS dataset. Both methods spend time on preprocessing steps that will be amortized over a large number of queries. The running time of APPLES*\**, as expected, increases linearly with *k >* 200 and grow more slowly for *k <* 200 (due to the preprocessing) (Fig. 4a). The patterns of running time of EPA-ng are surprising. Increasing *k* from 1 to 768 *decreases* the running time instead of increasing it. While the exact reasons for the reductions are not clear to us, we note that EPA-ng developers have taken great care to implement various optimizations, especially for utilizing multiple cores. The current version of APPLES*\**, in contrast, simply treats all queries as independent. Only with *k >* 768 we start to see increasing running times for EPA-ng. With *k >* 3072, EPA-ng seems to grow in running with *k*^0.73^; however, we suspect further increasing *k* would increase the exponent (asymptotic running time cannot theoretically be less than *O*(*k*) as all queries need to be read). Aside from the fluctuation due to small sample size for small *k* values, the number of queries do not seem to affect the accuracy for either method (Fig. 4b).

**Fig. 4.**
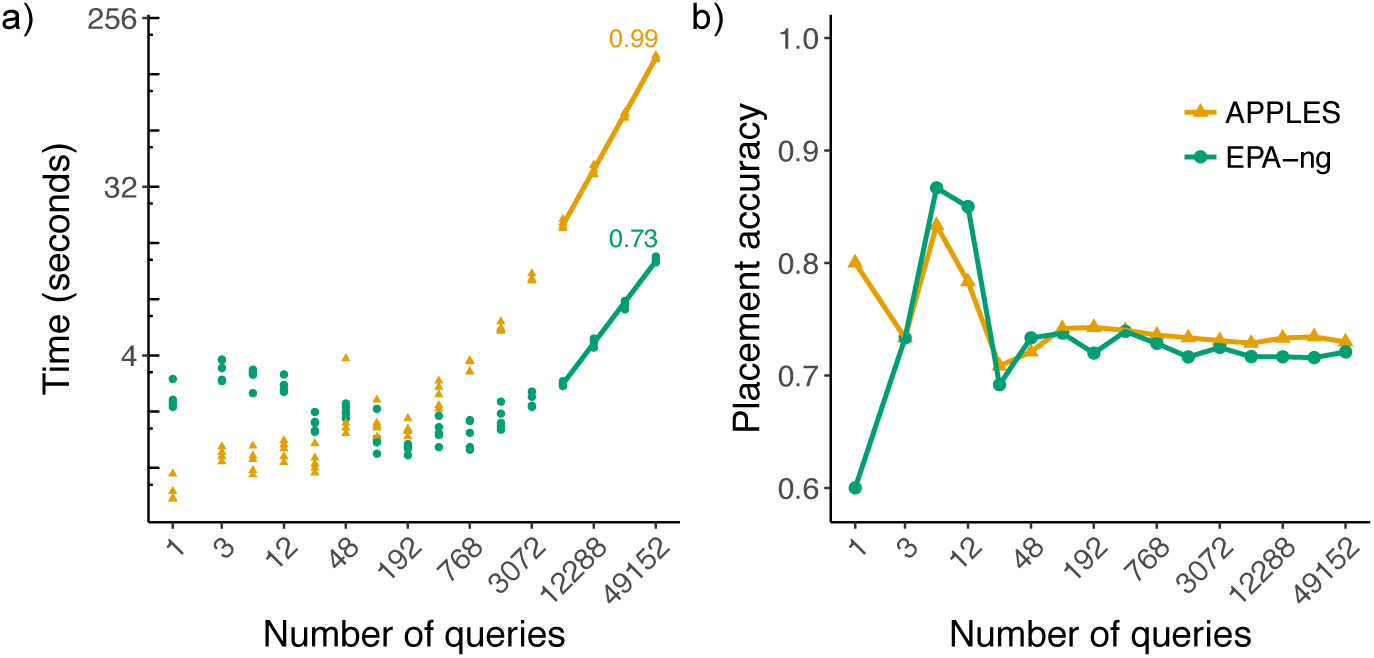
Scalability with respect to the number of queries. We show wall-clock running time (a) and accuracy with respect to increased numbers of queries (*k*) in one execution given 28 CPU cores on a Intel Xeon E5 CPU with 64 GB of memory. We fit a line to running times in log-log scale only for *k* ≥ 6144 because otherwise, the preprocessing time would distort estimates (note: *t* = *n*^*a*^ + *b* ⇏log *t* = *a* log *n* + *E* except in approximation if *b* ≪*t*).

#### Comparing parameters of APPLES

Comparing five models of sequence evolution available in APPLES*\**, we see similar patterns of accuracy across all models despite their varying complexity, ranging from 0 to 12 parameters (Fig. S3). Since the JC69 model is parameter-free and results in similar accuracy to others, we have used it as the default. Next, we ask whether imposing the constraint to disallow negative branch lengths improves the accuracy. The answer depends on the optimization strategy. Forcing non-negative lengths marginally increases the accuracy for MLSE but dramatically reduces the accuracy for ME (Fig. S4ab). Thus, we always impose non-negative constraints on MLSE but never for ME. Likewise, our Hybrid method includes the constraint for the first MLSE step but not for the following ME step (Fig. S4c).

The next parameter to choose is the weighting scheme. Among the three methods available in APPLES, the best accuracy belongs to the FM scheme closely followed by the BE (Fig. S5). The OLS scheme, which does not penalize long distances, performs substantially worse than FM and BE. Thus, the most aggressive form of weighting (FM) results in the best accuracy. Fixing the weighting scheme to FM and comparing the three optimization strategies (MLSE, ME, and Hybrid), the MLSE approach has the best accuracy (Fig. 2), finding the correct placement 84% of the time (mean error: 0.18), and ME has the lowest accuracy, finding the best placement in only 67% of cases (mean error: 0.70). The Hybrid approach is between the two (mean error: 0.34) and fails to outperform MLSE on this dataset. However, when we restrict the RNASim backbone trees to only 20 leaves, we observe that Hybrid can have the best accuracy (Fig. S6).

### Comparison to ML with Alignment Error

We next test the impact of alignment errors. On RNASim-AE dataset with *n* = 200, 000 sequences, we observe 80% placement accuracy using SEPP+APPLES*\** (Fig. 5), only 7% less than placing on the true backbone alignment with the true alignment of the query to the backbone. Similarly, on the *n* = 10, 000 backbone, placement accuracy drops from 83% on the true alignment to 80% using SEPP+APPLES*\**, and from 81% to 78% using SEPP+EPA-ng. Despite the relatively small drops on placement accuracy, the impact of alignment error on delta error can be more pronounced (Fig. 5). The mean delta error for *n* = 10, 000 and *n* = 200, 000 goes up from 0.33 and 0.25 edges, respectively, to 0.48 and 0.55 edges.

**Fig. 5.**
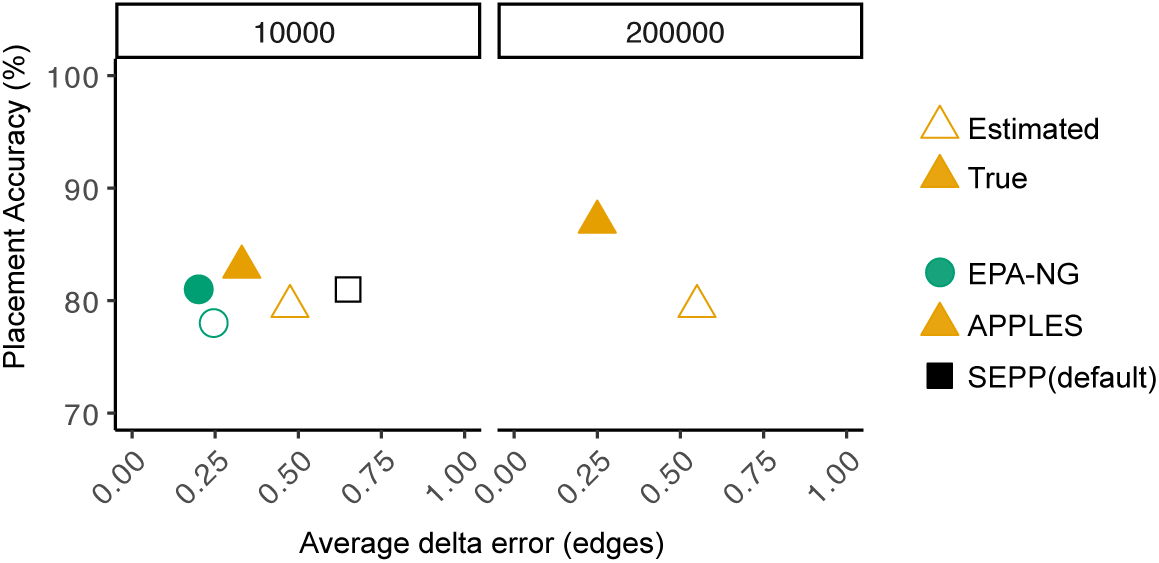
Impact of alignment error on placement accuracy. Left and right panels show placement accuracy and mean delta error of leave-many-out experiments for backbone size of 10, 000 and 200, 000 (200 queries each).

Recall that SEPP+APPLES*\** eliminates the need for decomposing the backbone tree into smaller placement subtrees, as default SEPP must do to deal with memory requirements of pplacer (which it internally uses). Comparison of default SEPP and SEPP+APPLES*\** shows that incorporating APPLES*\** inside SEPP reduces the mean delta error from 0.65 to 0.48, while it hurts placement accuracy by 1% (81% vs 80%). To summarize, highly accurate placement is possible even with estimated alignments with backbones of size up to 200, 000 using the divide-and-conquer methods PASTA and SEPP for estimating alignments of the backbone and query, respectively.

## Discussion

We introduced APPLES: a new method for adding query species onto large backbone trees using both unassembled genome-skims and aligned data. We now provide further observations on our results and on distance-based placement.

### Further observations on the results

The accuracy of APPLES was very close to ML in most settings where we could run ML; the accuracy advantages of ML were particularly small for the RNASim dataset where both methods face model misspecification. As expected by the substantial evidence from the literature (Hillis *et al.*, 2003; Zwickl and Hillis, 2002), improved taxon sampling increased the accuracy of placement. Thus, overall, the best accuracy on RNASim dataset was obtained by APPLES*\** run on the full reference dataset, further motivating its use when large backbones are available. Despite many strides made in terms of scalability by the new method EPA-ng, ML methods still have to restrict their backbone to at most several thousand species given reasonable amounts of memory (up to 64GB in our case). We also note that it is possible to follow the APPLES*\** placement with a round of ML placement on smaller trees, but the small differences in accuracy of ML and APPLES*\** on smaller trees did not give us compelling reasons to try such hybrid approaches.

APPLES was an order of magnitude or more faster and less memory-hungry than ML tools (pplacer and EPA-ng) for single query runs. However, for placing large numbers of queries (e.g., as found in metagenomic datasets) on a relatively small sized backbone (*n* = 500), EPA-ng had an advantage thanks to heuristics it employs and carefully engineered code. Advantages in memory consumption and scalability to large backbone trees remain for APPLES*\** regardless of the number of queries. The python APPLES code is not optimized nearly as much as EPA-ng and can also benefit from some of the heuristic techniques used by EPA-ng. We plan for the future versions of the code to focus on improved scalability as the number of queries increases.

By incorporating APPLES inside SEPP, we were able to create a method that can do both alignment and placement on very large backbones with reasonable computational requirements and high accuracy. We observed relatively small reductions in accuracy as a result of alignment error, a pattern that we find remarkable given the size of the tree and the amount of error in the estimated alignment. The default SEPP method deals with large backbone trees by dividing the backbone tree into “placement” subtrees and choosing which subtree to place on using bit-scores produced by HMMs trained on subsets of sequences. The original paper had shown that the best accuracy is obtained with the largest possible backbone subtrees given computational limitations. APPLES now enables us to eliminate the need for decomposition into placement subsets, and in doing so, reduces placement error.

In our analyses, we observed no advantage in using models more complex than JC69+Γ for distance calculation inside APPLES. However, these results may be due to our pairwise estimation of model parameters (e.g., base compositions). More complex models may perform better if we instead estimate model parameters on the backbone alignment/tree and reuse the parameters for queries (or simultaneously among all queries and the reference sequences). Simultaneous estimation of distances has many advantages over using independent distances for the *de novo* case (Tamura *et al.*, 2004; Xia, 2009); these results give us hope that using simultaneous distances inside APPLES can further improve its accuracy.

Branch lengths of our backbone trees were computed using the same distance model as the one used for computing the distance of the query to backbone species. Using consistent models for the query and for the backbone branch lengths is essential for obtaining good accuracy (see Fig. S7 for evidence). Thus, in addition to having a large backbone tree at hand, we need to ensure that branch lengths are computed using the right model. Fortunately, FastTree-2 can compute both topologies and branch lengths on large trees in a scalable fashion, without a need for quadratic time/memory computation of distance matrices (Price *et al.*, 2010).

### Observations on distance-based placement

Phylogenetic insertion using the ME criterion has been previously studied for the purpose of creating an algorithm for greedy minimum evolution (GME). Desper and Gascuel 2002 have designed a method that given the tree *T* can update it to get a tree with *n* + 1 leaves in Θ(*n*) after precomputation of a data-structure that gives the average *sequence* distances between all adjacent clusters in *T*. The formulation by Desper and Gascuel 2002 has a subtle but consequential difference from our ME placement. Their algorithm does not compute branch lenghts for inserted sequence (e.g., *x*_1_ and *x*_2_). It is able to compute the optimal placement topology *without* knowing branch lengths of the backbone tree. Instead, it relies on pairwise distances among backbone *sequences* (Δ), which are precomputed and saved in the data-structure mentioned before. In the context of the greedy algorithm for tree inference, in each iteration, the data structure can be updated in Θ(*n*), which does not impact the overall running time of the algorithm. However, if we were to start with a tree of *n* leaves, computing this structure from scratch would still require Θ(*n*^2^). Thus, computing the placement for a new query would need quadratic time, unless if the Θ(*n*^2^) precomputation is allowed to be amortized over Ω(*n*) queries. Our formulation, in contrast, uses branch lengths of the backbone tree (which is assumed fixed) and thus never uses pairwise distances among the backbone sequences. Thus, using tree distances is what allows us to develop a linear time algorithm. Finally, we note that in our experimental analyses, we were not able to test the distance-based algorithm of Desper and Gascuel (2002) because it is available only as part of a the greedy algorithm inside FastME but is not available as a stand-alone feature to place on a given tree.

We emphasize that our results in assembly-free tests do not advocate for the use of assembly-free methods when assemblies are available. Moreover, we have no evidence that assembly-free methods are effective in inferring deep branches of a phylogeny. Instead, our results show that assembly-free phylogenetic placement is effective in sample identification where assembly is not possible due to low coverage. In assembly-free analyses, we used Skmer to get distances because alternative alignment-free methods of estimating distance generally either require assemblies (e.g., Haubold, 2014; Leimeister and Morgenstern, 2014; Leimeister *et al.*, 2017) or higher coverage than Skmer (e.g., Benoit *et al.*, 2016; Yi and Jin, 2013; Ondov *et al.*, 2016); however, combining APPLES with other alignment-free methods can be attempted in future (finding the best way of computing distances without assemblies was not our focus). Moreover, the Skmer paper has described a trick that can be used to compute log-det distances from genome-skims. Future studies should test whether using that trick and using GTR instead of JC69 improves accuracy.

## Acknowledgments

This work was supported by the National Science Foundation (NSF) grant IIS-1565862 and National Institutes of Health (NIH) subaward 5P30AI027767-28 to M.B. and S.M., and NSF grant NSF-1815485 to M.B., S.S., and S.M. Computations were performed on the San Diego Supercomputer Center (SDSC) through XSEDE allocations, which is supported by the NSF grant ACI-1053575.

## Appendix

### APPENDIX A. Proofs and Derivations

Recall the following notations.

- For any node *u* and exponents *a* ∈ 𝕫 and *b* ∈ ℕ^+^, let
  — 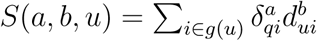
  — 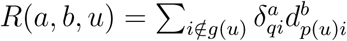 defined for *u* ∊ *V* \ {1}
- For *b* = 0, let 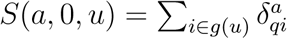 and let *S′*(*a, u*) be a shorthand for *S*(*a*, 0, *u*). Similarly, let 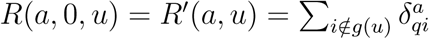.

#### Proof of Lemma 2

*Proof* Recall the dynamic programming recursions of Equations 3 and 4:

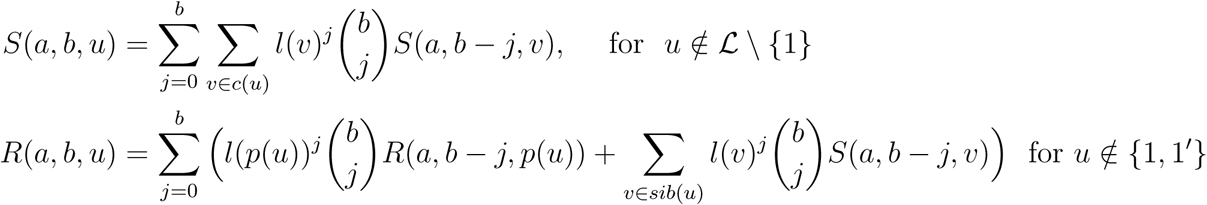

Since *u* is not a leaf, for each leaf *i* ∈ *g*(*u*), there exists a *v* ∈ *c*(*u*) such that the directed path from *u* to *i* passes through *v*. Therefore every leaf *i* can be grouped under its corresponding *v*.

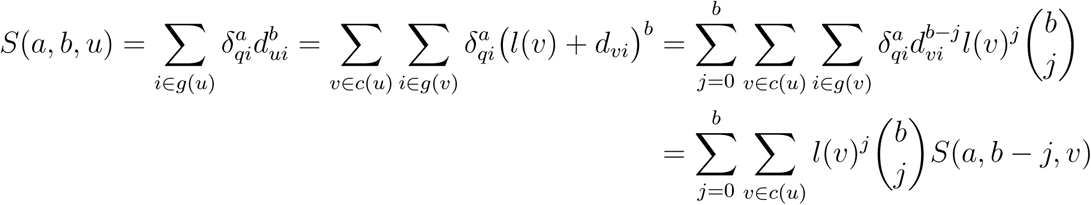

Similarly, given the condition *u* ≠ 1, for each leaf *i* ∉*g*(*u*), either (1) there exists *v* ∈ *sib*(*u*) such that the directed path from *p*(*u*) to *i* passes through *v*, or (2) undirected path between *i* and *p*(*u*) passes through *p*(*p*(*u*)).

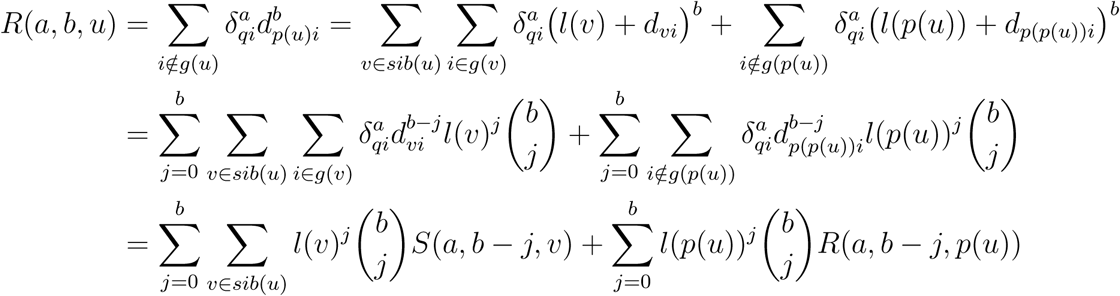

Boundary conditions follow from definitions. For *u* ∉*L \ {*1}, since *d*_*ii*_ = 0, we have *S*(*a, b, u*) = 0 and it’s trivial to see 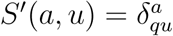. For *R*(,,) recursions, the boundary case happens at the unique child of the root, which we denote as 1′. Based on the definition, since the only *i* ∉ *g*(1^*′*^) is 1, and 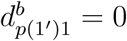, we trivially have *R*(*a, b*, 1^*′*^) = 0. For *b* = 0,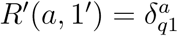.

A post-order traversal on *T\** can compute *S*(*a, b, u*), and a subsequent pre-order traversal can compute *R*(*a, b, u*), both in constant time and using constant memory per node. Recall that *a* and *b* are both no more than *k*, which is a constant. Thus, time and memory complexity of this dynamic programming is Θ(*bn*), which translates to Θ(*n*) in least squares setting, where *b* ≤ 2. □

#### Proof of Lemma 3

Recall 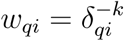 and that Equation 2:

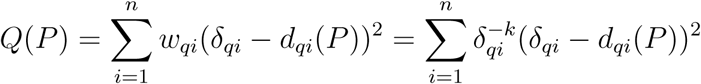

*Proof*. Equation 2 can be re-written as:

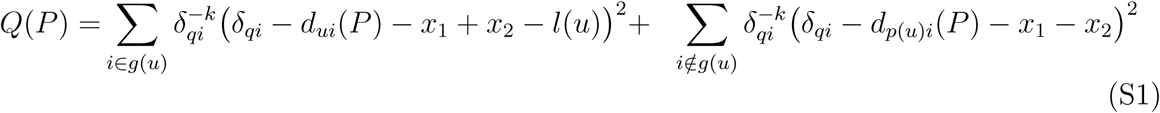

By simple rearrangement of the terms, we can rewrite Equation S1 as follows.

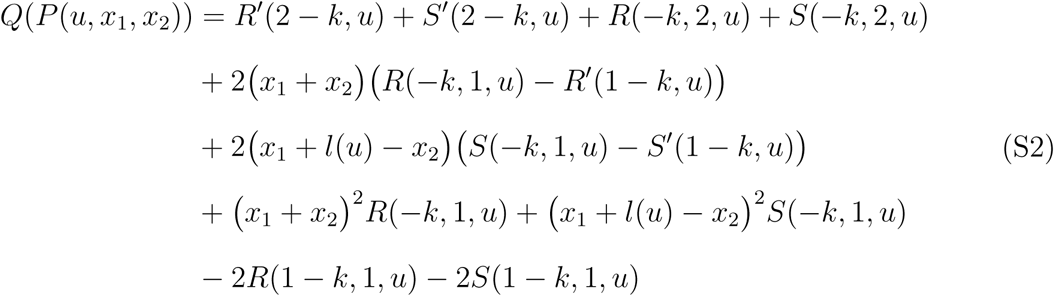

Note that computing *Q*(*P* (*u, x*_1_, *x*_2_)) requires only *S*(,, *u*) and *R*(,, *u*) values and *l*(*u*). Thus, computing *Q*(*P*) requires only computing *S*(*a, b, u*) and *R*(*a, b, u*) values for −*k* ≤*a* ≤ 2 − *k* and 0 ≤*b* ≤ 2.

□

#### Proof of Lemma 4

Recall definitions

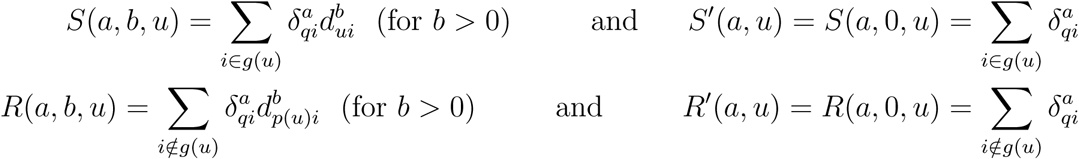

and recall Eq. S1:

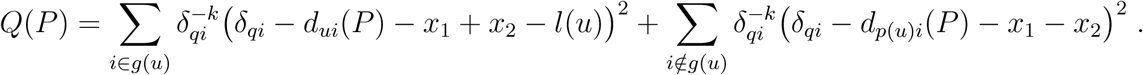

*Proof.* We take the derivative of Eq. S1 with respect to *x*_1_ and set it equal to zero:

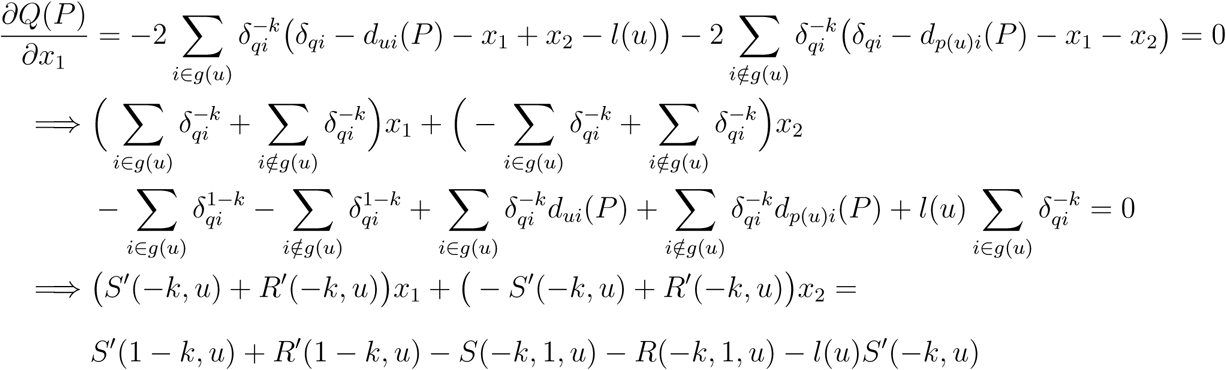

Similarly,

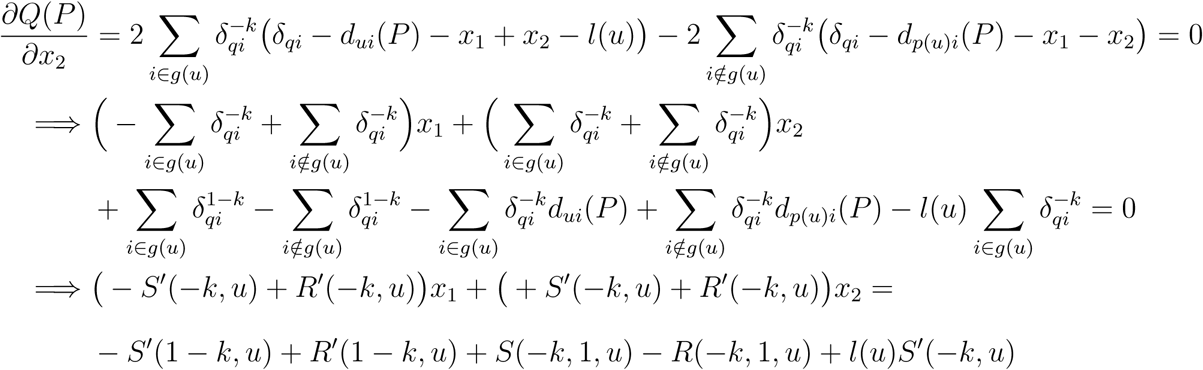

These two linear equations have a unique solution for the pair *x*_1_, *x*_2_ if and only if the following matrix has the full rank:

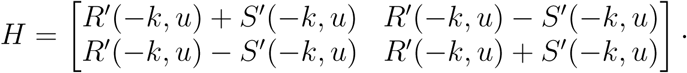

Determinant of *H* is det(*H*) = 4*R′*(−*k, u*)*S′*(−*k, u*). Assuming that *δ*_*qi*_ *>* 0 for all *i* ∈ *ℒ*, both *R′*(−*k, u*) *>* 0 and *S′*(−*k, u*) *>* 0 hold. Therefore, *H* has the full rank. However, *δ*_*qi*_ = 0 for *q* ≠ *i* can be encountered on real data, especially for low divergence times, low evolutionary rates, or short sequences. In this case, APPLES is designed to place *q* on the pendant edge of *i* with *x*_1_ = 0 and *x*_2_ = *l*(*i*). In case there are multiple leaves *i* that satisfy *δ*_*qi*_ = 0 for *q* ≠ *i*, we pick one of them arbitrarily. □

#### Proof of Theorem 1

*Proof.* First, using two traversals of the tree, we compute all the *S*(*a, b, u*) and *R*(*a, b, u*) values by Lemma 2. To find the optimal placement edge, we first optimize *Q*(*P* (*u, x*_1_, *x*_2_)) for all *u* ∈ *V* \ {1}. By Lemma 4, this task requires only constant time after the precompuations. Then, for each node, we compute *Q*(*P* (*u, x*_1_, *x*_2_)) in constant time for the optimal *u, x*_1_, *x*_2_ by Lemma 3. Thus, each node is processed in linear time and the whole optimization requires linear time. Note that the system of equations (shown in Lemma 4) will not have a solution iff *δ*_*qi*_ ≤ 0 for some *i*; if there is *δ*_*qi*_ = 0, we make *q* sister to *i*, breaking ties arbitrarily. □

#### Proof of Lemma 5

*Proof.* Eigenvalues of the Hessian matrix of *Q*(*P* (*u, x*_1_, *x*_2_)) are 2*R′*(−*k, u*) and 2*S′*(−*k, u*), which are both non-negative since *δ*_*qi*_ ≥ 0 for *i* ∈ *ℒ*. Thus, the Hessian matrix is positive semidefinite and therefore *P* (*u, x*_1_, *x*_2_) is a convex function of *x*_1_ and *x*_2_. □

### APPENDIX B. Supplementary Figures

**Fig. S1.**
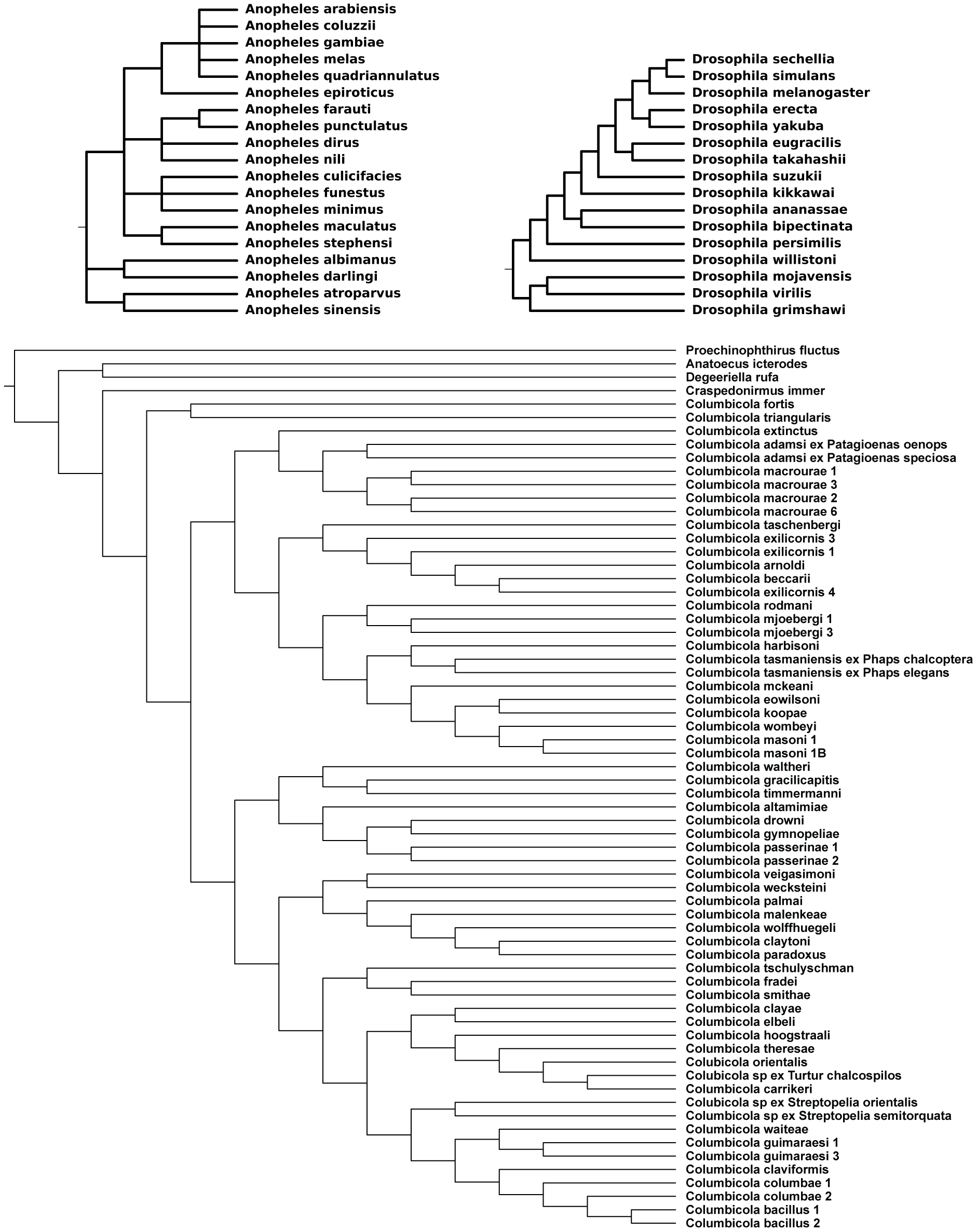
The reference biological trees obtained from Open Tree of Life (Drosophila and Anopheles) and from Boyd *et al.* (2017) (Columbicola).

**Fig. S2.**
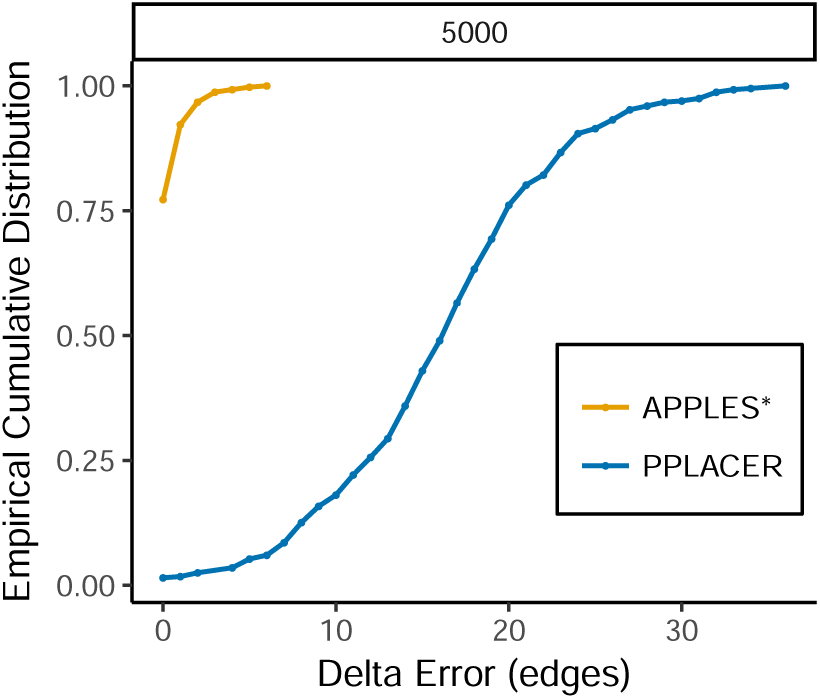
APPLES versus pplacer on 5,000 backbone trees. The empirical cumulative distribution of the delta error is shown. We compare pplacer and APPLES*\** on RNASim-VS dataset with 5000 leaves. Distributions is over 551 cases where pplacer could run for the panel.

**Fig. S3.**
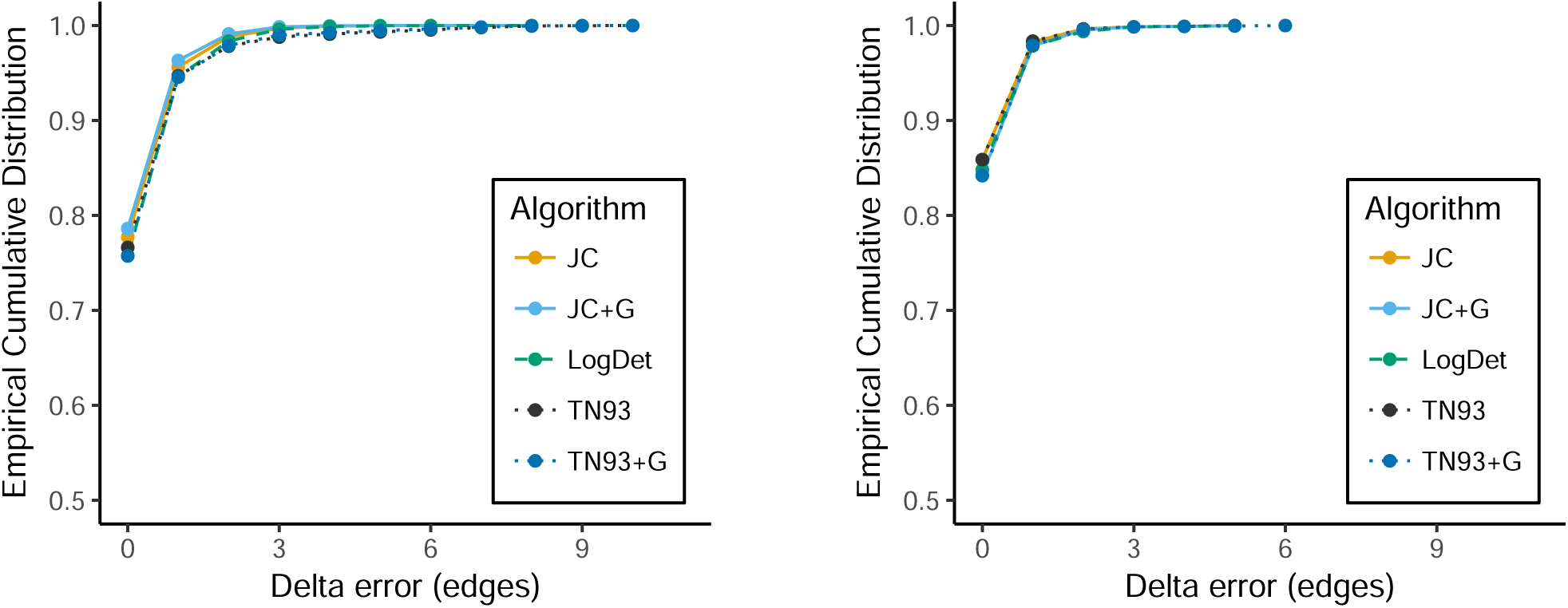
Comparing various models of DNA evolution. For the GTR (a) and RNASim-heterogeneous (b) datasets, we show the delta error (edges) of APPLES* run with five distance matrices calculated based on different models of DNA evolution. All model parameters are estimated per pair of sequences. The five models have similar accuracy.

**Fig. S4.**
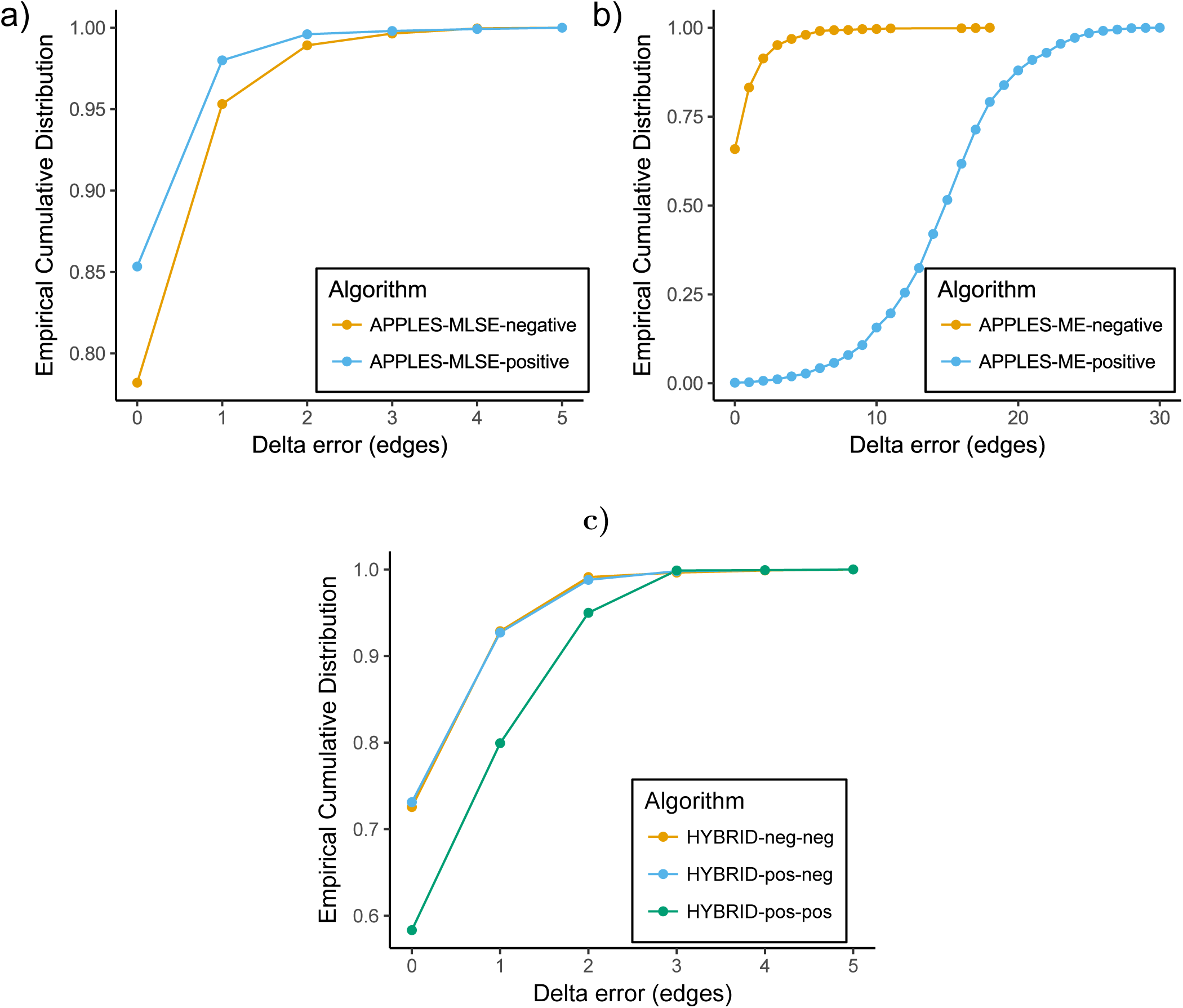
(a,b) The effect of imposing positivity constraint on error. We show the error of (a) APPLES-MLSE and (b) APPLES-ME run both with and without enforcement of non-negative branch lengths on RNASim heterogeneous dataset. Accuracy improves substantially for MLSE whereas it reduces drastically for ME. **(c) The effect of imposing positivity constraint on accuracy on Hybrid.** The HYBRID approach does not benefit from imposing positivity constraint on its second (ME) stage.

**Fig. S5.**
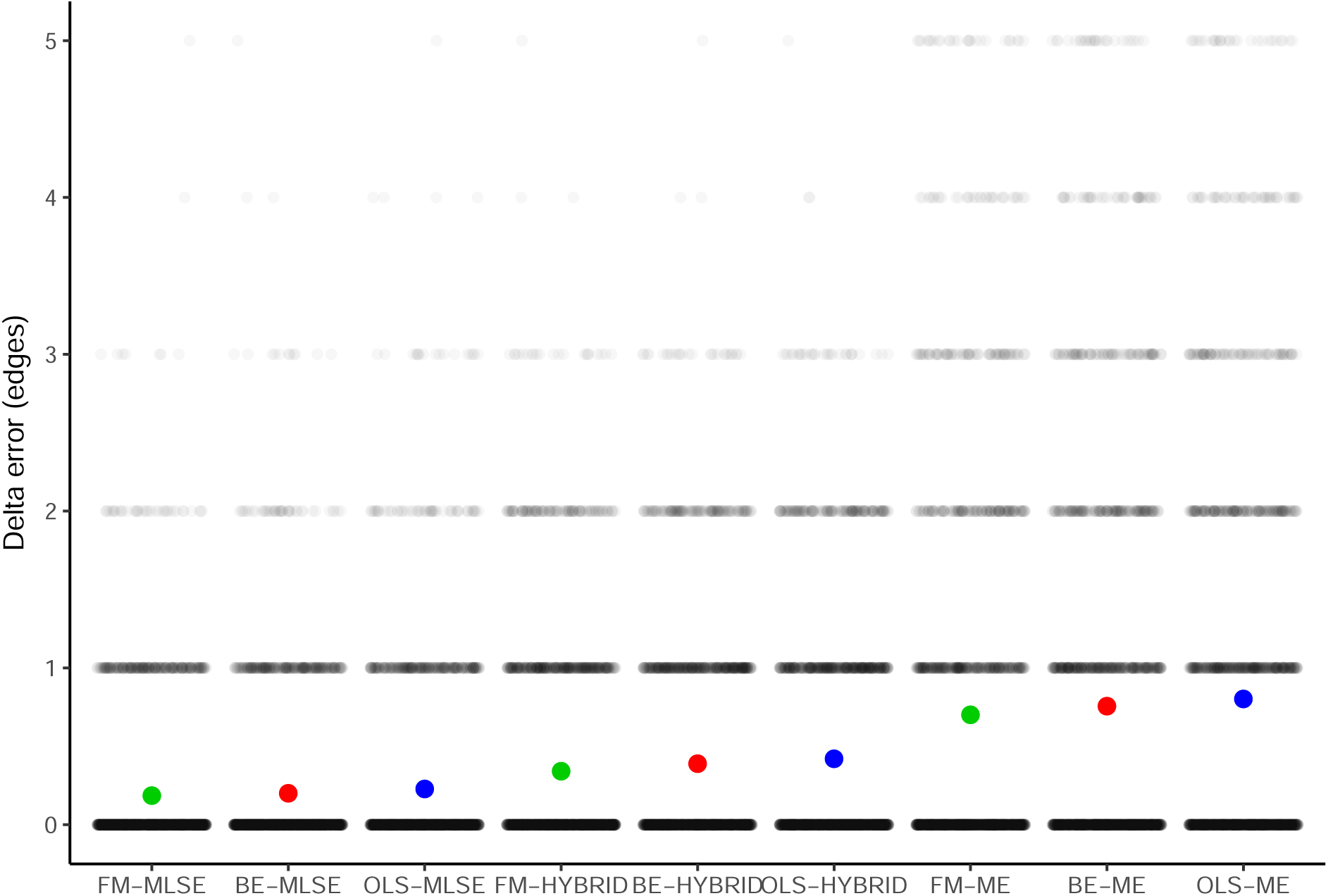
Comparing APPLES versions. For the RNASim dataset (without controlling for the diameter), we show the delta error (edges) of APPLES run with three options for weighting: FM (green), BE (red), and OLS (blue), and three options for selection strategy (MLSE, ME, and Hybrid). For each method, the mean (colored circle) and standard errors (lines; too small to see) are shown over 2500 data points, each shown as dots. Some of the methods occasionally have error above 5 branches, but for better resolution, we cap the y-axis at 5.

**Fig. S6.**
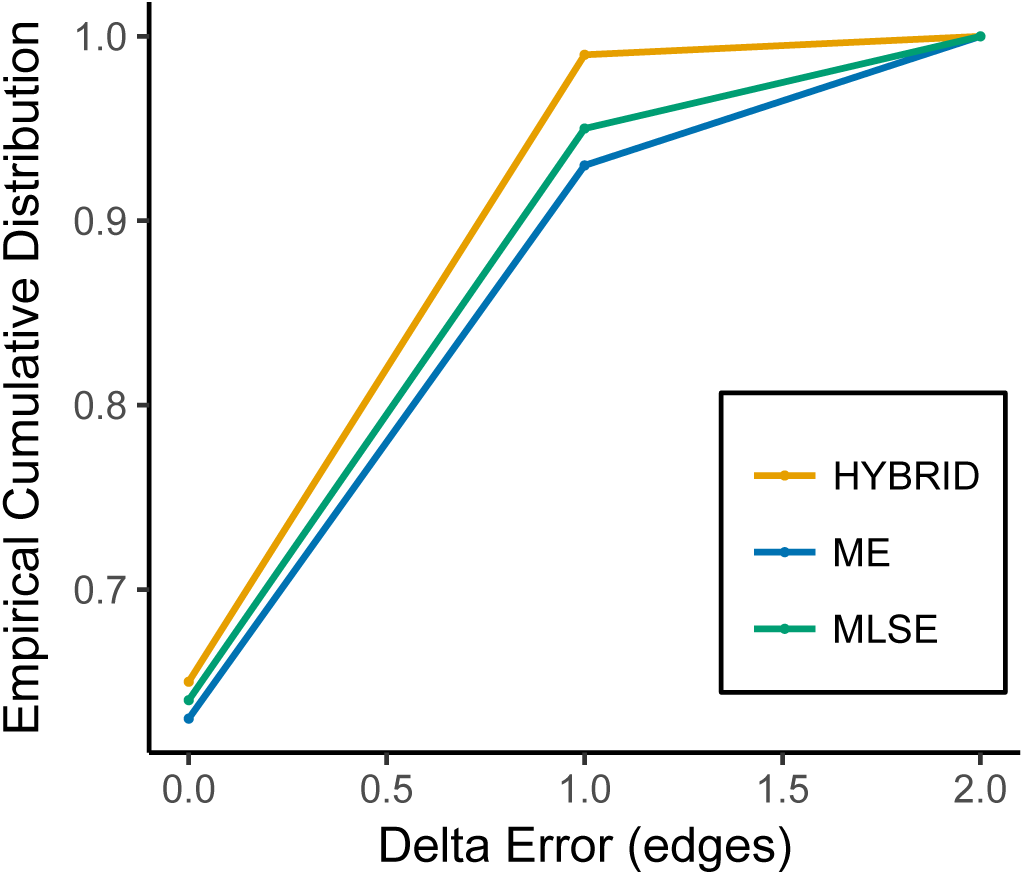
APPLES-HYBRID has higher accuracy on sparse RNAsim dataset. On the RNAsim dataset, we chose 20 sequences randomly from the larger RNAsim-heterogeneous dataset; here, APPLES-HYBRID has higher accuracy than APPLES* (MLSE).

**Fig. S7.**
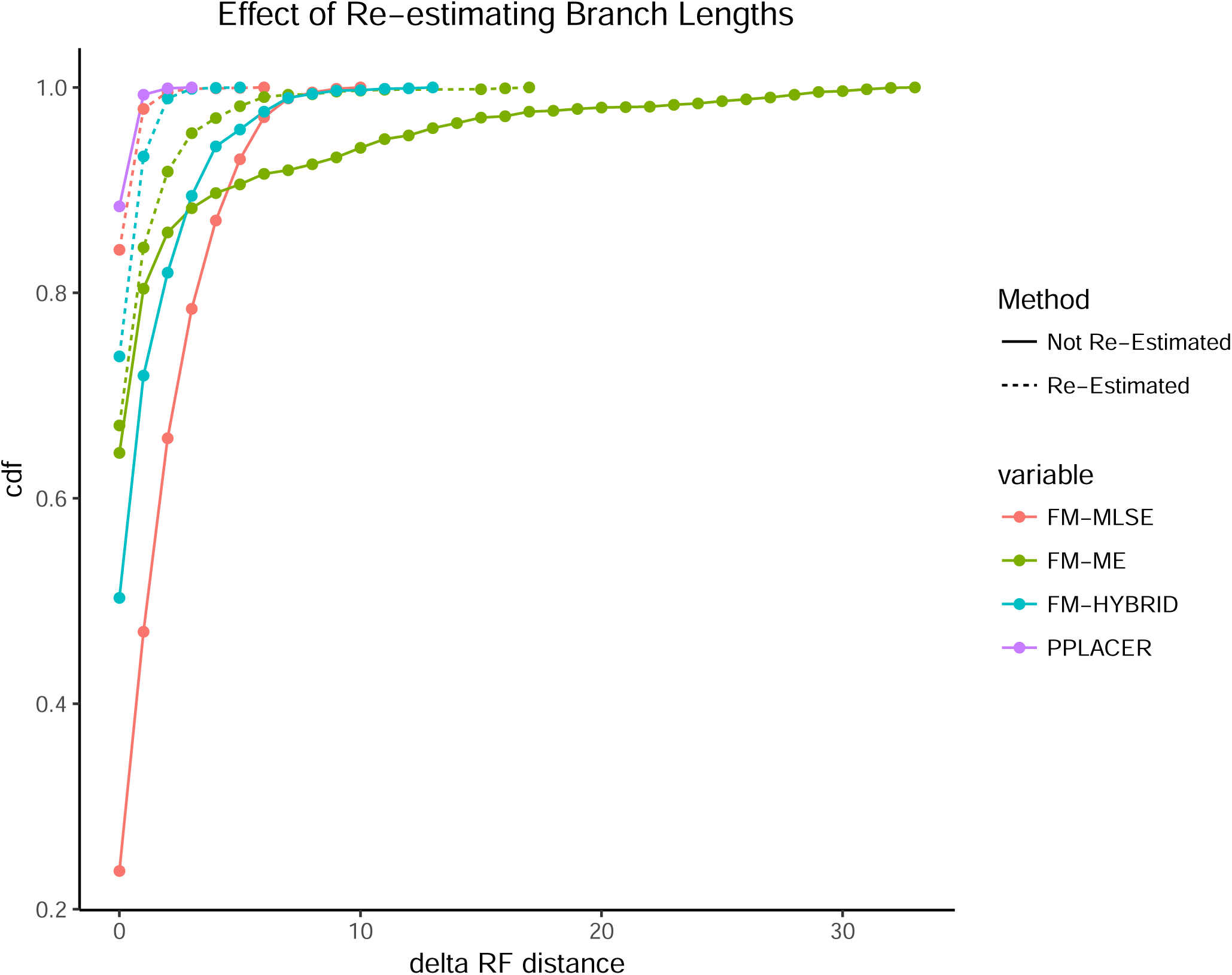
The effect of reestimating branch lengths of the backbone tree on accuracy. We show the accuracy of pplacer and APPLES-FM with its three optimization criteria. APPLES is run both with (dotted) and without (solid) re-estimating branch lengths in the backbone tree using the same model (here, TN93+Γ) used for computing distances of query sequences to backbone sequences. FastME* is used to re-estimate branch lengths. Accuracy improves dramatically by recomputing backbone branch lengths using the same model. The case labeled “Not re-estimated” uses branch lengths produced using RAxML under the GTR+Γ model.

### APPENDIX C. Supplementary Tables

**Table S1.** GenBank accession numbers and URLs for the dataset of 22 Anopheles genomes

| Species | GenBank assembly accession | URL |
| --- | --- | --- |
| *Anopheles albimanus* | GCA_000349125.1 | http://www.insect-genome.com/data/genome_download/Anopheles_albimanus/Anopheles_albimanus_genomic.fasta.gz |
| *Anopheles arabiensis* | GCA_000349185.1 | http://www.insect-genome.com/data/genome_download/Anopheles_arabiensis/Anopheles_arabiensis_genomic.fasta.gz |
| *Anopheles atroparvus* | GCA_000473505.1 | http://www.insect-genome.com/data/genome_download/Anopheles_atroparvus/Anopheles_atroparvus_genomic.fasta.gz |
| *Anopheles christyi* | GCA_000349165.1 | http://www.insect-genome.com/data/genome_download/Anopheles_christyi/Anopheles_christyi_genomic.fasta.gz |
| *Anopheles coluzzii* | - | http://www.insect-genome.com/data/genome_download/Anopheles_coluzzii/Anopheles_coluzzii_genomic.fasta.gz |
| *Anopheles culicifacies* | GCA_000473375.1 | http://www.insect-genome.com/data/genome_download/Anopheles_culicifacies/Anopheles_culicifacies_genomic.fasta.gz |
| *Anopheles darlingi* | GCA_000211455.3 | http://www.insect-genome.com/data/genome_download/Anopheles_darlingi/Anopheles_darlingi_genomic.fasta.gz |
| *Anopheles dirus* | GCA_000349145.1 | http://www.insect-genome.com/data/genome_download/Anopheles_dirus/Anopheles_dirus_genomic.fasta.gz |
| *Anopheles epiroticus* | GCA_000349105.1 | http://www.insect-genome.com/data/genome_download/Anopheles_epiroticus/Anopheles_epiroticus_genomic.fasta.gz |
| *Anopheles farauti* | GCA_000956265.1 | http://www.insect-genome.com/data/genome_download/Anopheles_farauti/Anopheles_farauti_genomic.fasta.gz |
| *Anopheles funestus* | GCA_000349085.1 | http://www.insect-genome.com/data/genome_download/Anopheles_funestus/Anopheles_funestus_genomic.fasta.gz |
| *Anopheles gambiae* | GCA_000150785.1 | http://www.insect-genome.com/data/genome_download/Anopheles_gambiae/Anopheles_gambiae_genomic.fasta.gz |
| *Anopheles koliensis* | GCA_000956275.1 | http://www.insect-genome.com/data/genome_download/Anopheles_koliensis/Anopheles_koliensis_genomic.fasta.gz |
| *Anopheles maculatus* | GCA_000473185.1 | http://www.insect-genome.com/data/genome_download/Anopheles_maculatus/Anopheles_maculatus_genomic.fasta.gz |
| *Anopheles melas* | GCA_000473525.2 | http://www.insect-genome.com/data/genome_download/Anopheles_melas/Anopheles_melas_genomic.fasta.gz |
| *Anopheles merus* | GCA_000473845.2 | http://www.insect-genome.com/data/genome_download/Anopheles_merus/Anopheles_merus_genomic.fasta.gz |
| *Anopheles minimus* | GCA_000349025.1 | http://www.insect-genome.com/data/genome_download/Anopheles_minimus/Anopheles_minimus_genomic.fasta.gz |
| *Anopheles nili* | GCA_000439205.1 | http://www.insect-genome.com/data/genome_download/Anopheles_nili/Anopheles_nili_genomic.fasta.gz |
| *Anopheles punctulatus* | GCA_000956255.1 | http://www.insect-genome.com/data/genome_download/Anopheles_punctulatus/Anopheles_punctulatus_genomic.fasta.gz |
| *Anopheles quadriannulatus* | GCA_000349065.1 | http://www.insect-genome.com/data/genome_download/Anopheles_quadriannulatus/Anopheles_quadriannulatus_genomic.fasta.gz |
| *Anopheles sinensis* | GCA_000441895.2 | http://www.insect-genome.com/data/genome_download/Anopheles_sinensis/Anopheles_sinensis_genomic.fasta.gz |
| *Anopheles stephensi* | GCA_000300775.2 | http://www.insect-genome.com/data/genome_download/Anopheles_stephensi/Anopheles_stephensi_genomic.fasta.gz |

**Table S2.**
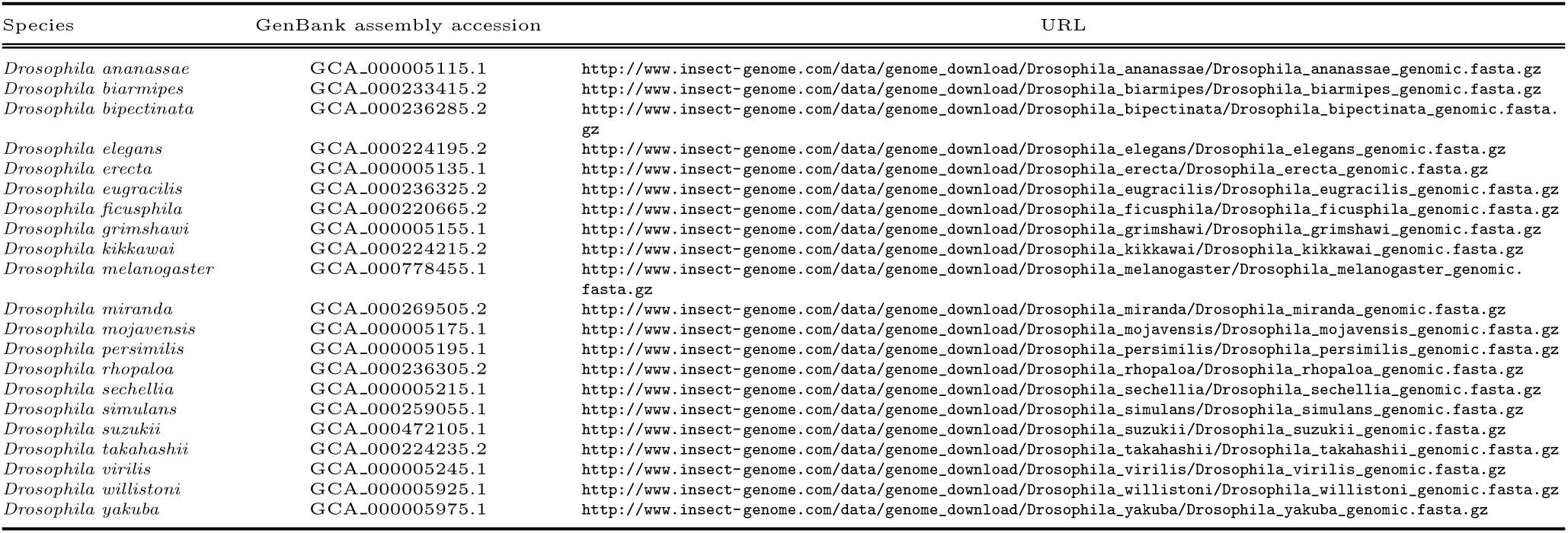
GenBank accession numbers and URLs for the dataset of 21 Drosophila genomes

**Table S3.**
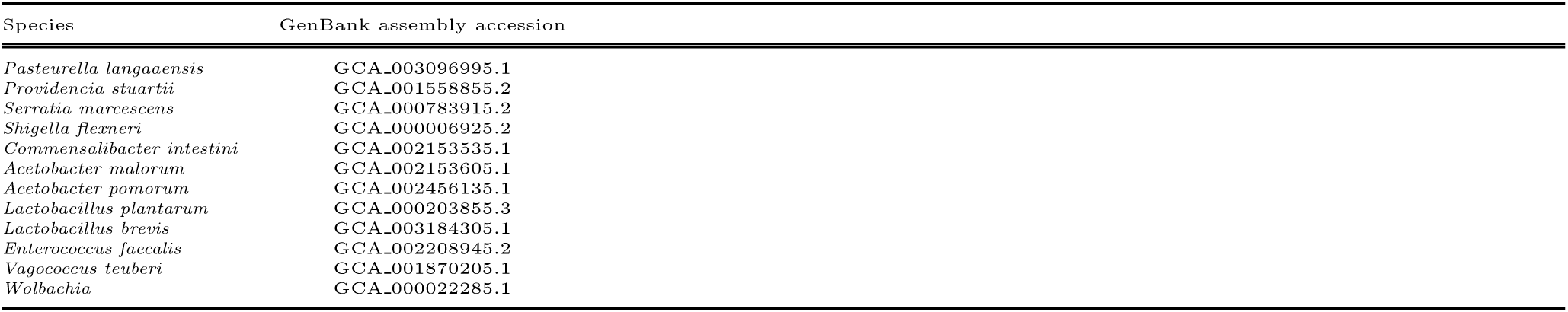
GenBank accession numbers of microbial species used in contamination removal.

**Table S4.**
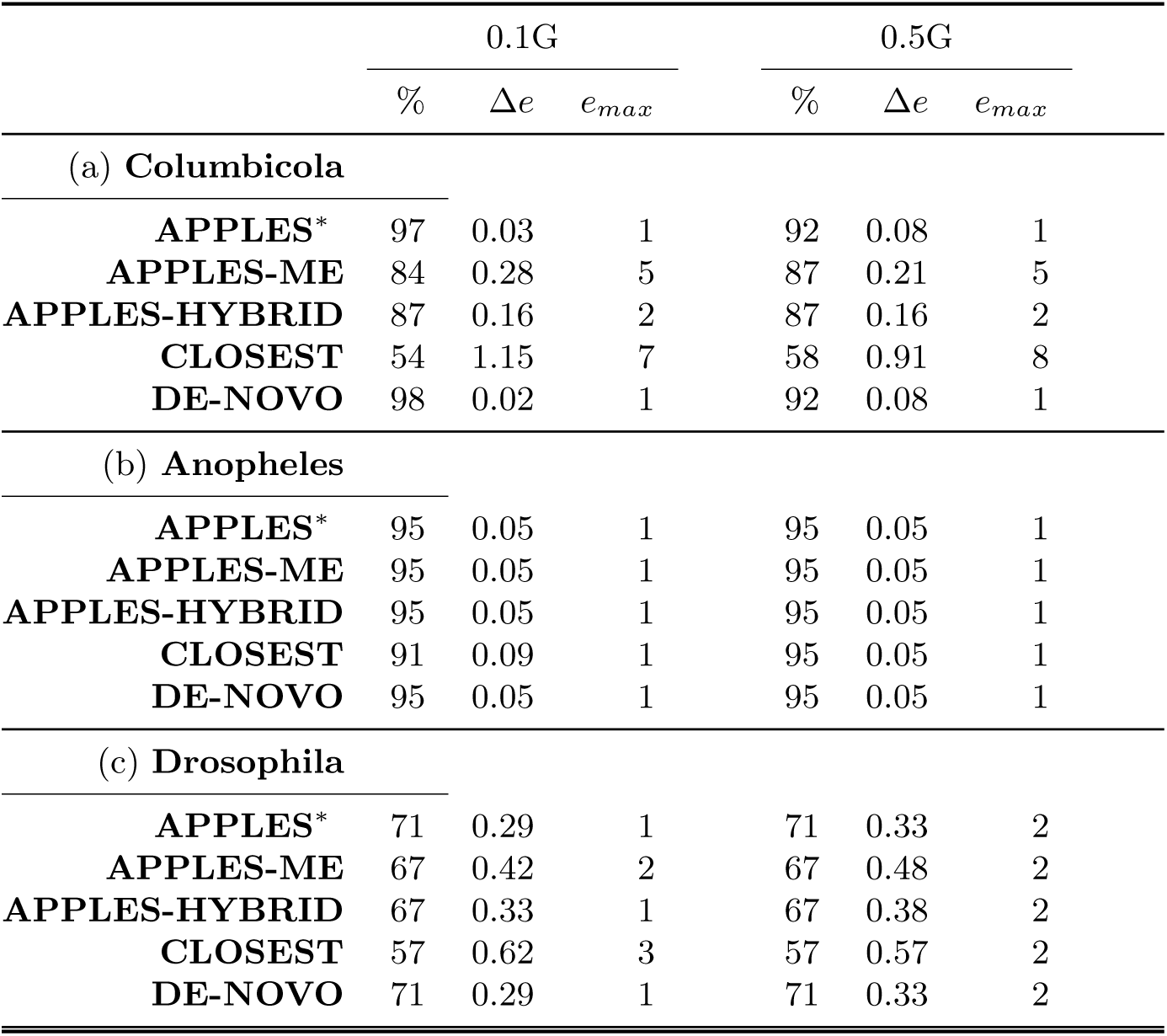
Assembly-free placement of genome-skims. We show the percentage of correct placements (those that do not increase Δ*e*), average delta error (Δ*e*), and maximum delta error (*e*_*max*_) for APPLES, assignment to the CLOSEST species, and the placement to the position in the backbone (DE-NOVO) over the 61 (a), 22 (b), and 21 (c) placements. Results are shown for skims with 0.1 and 0.5Gbp of reads. Delta error is the increase in the number missing branches between the reference tree and the backbone tree after placing each query.

|  | 0.1G |  |  | 0.5G |  |  |
| --- | --- | --- | --- | --- | --- | --- |
| | % | $\Delta e$ | $e_{max}$ | % | $\Delta e$ | $e_{max}$ |
| (a) **Columbicola** |  |  |  |  |  |  |
| APPLES* | 97 | 0.03 | 1 | 92 | 0.08 | 1 |
| APPLES-ME | 84 | 0.28 | 5 | 87 | 0.21 | 5 |
| APPLES-HYBRID | 87 | 0.16 | 2 | 87 | 0.16 | 2 |
| CLOSEST | 54 | 1.15 | 7 | 58 | 0.91 | 8 |
| DE-NOVO | 98 | 0.02 | 1 | 92 | 0.08 | 1 |
| (b) **Anopheles** |  |  |  |  |  |  |
| APPLES* | 95 | 0.05 | 1 | 95 | 0.05 | 1 |
| APPLES-ME | 95 | 0.05 | 1 | 95 | 0.05 | 1 |
| APPLES-HYBRID | 95 | 0.05 | 1 | 95 | 0.05 | 1 |
| CLOSEST | 91 | 0.09 | 1 | 95 | 0.05 | 1 |
| DE-NOVO | 95 | 0.05 | 1 | 95 | 0.05 | 1 |
| (c) **Drosophila** |  |  |  |  |  |  |
| APPLES* | 71 | 0.29 | 1 | 71 | 0.33 | 2 |
| APPLES-ME | 67 | 0.42 | 2 | 67 | 0.48 | 2 |
| APPLES-HYBRID | 67 | 0.33 | 1 | 67 | 0.38 | 2 |
| CLOSEST | 57 | 0.62 | 3 | 57 | 0.57 | 2 |
| DE-NOVO | 71 | 0.29 | 1 | 71 | 0.33 | 2 |

### Appendix D. Commands

#### Sampling Clades

For sampling clades of size at most 250 from a tree “tree.nwk”, we used the TreeCluster package, available at https://github.com/niemasd/TreeCluster.

~~~
*#!/bin/bash*
python TreeCluster/TreeCluster.py -i 250 -o clusters.txt -t tree.nwk
-m count_max_clade
~~~

#### Computing Distance Matrices

APPLES version 1.2.0 can compute JC69 distances between nucleotide sequences. Version 1.1.0 internally uses FastME* (based on FastME version 2.1.6.1) which is available at https://github.com/balabanmetin/FastME-personal-copy. We computed distance matrices based on other models (e.g. TN93+Γ) using following FastME command:

~~~
*#!/bin/bash*
fastme -c -dT --gamma=${gamma} -i aln_dna.phy -O dist.mat -T 1
~~~

where ${*gamma*} is Γ the rate variable.

#### Backbone tree estimation

When multiple sequence alignment is available, we used the following RAxML command to compute backbone tree for all datasets except RNAsim-VS dataset. We used RAxML version 7.2.6

~~~
*#!/bin/bash*
raxmlHPC-PTHREADS -m GTRGAMMA -p 88 -n REF -s aln_dna.phy -T 6
~~~

For RNAsim-VS dataset, we used FastTreeMP version 2.1.10 for estimating backbone topology. We run FastTreeMP with the following command:

~~~
*#!/bin/bash*
FastTreeMP -nosupport -gtr -gamma -nt -log tree.log < aln_dna.fa > tree.nwk
~~~

For alignment free datasets such as Drosophila dataset, we computed backbone tree using FastME* (based on FastME version 2.1.6.1) which is available at https://github.com/balabanmetin/FastME-personal-copy. FastME* is run with the following command:

~~~
*#!/bin/bash*
fastme -i dist.mat -o tree.nwk -T 1
~~~

Note that we performed Jukes-Cantor correction on the distance matrix “dist.mat” before running FastME*.

#### Backbone tree branch length re-estimation

When multiple sequence alignment is available, we used FastME* to recompute backbone tree branch lengths for all datasets except RNAsim-VS dataset. We run FastME* with the following command:

~~~
*#!/bin/bash*
fastme -dJ -i aln_dna.phy -u RAxML_result.REF -o tree_me.nwk
~~~

For RNAsim-VS dataset, we used RAxML version 7.2.6 for re-estimating ML based branch lengths and used that tree for performing placements using pplacer. RAxML is run with the following command:

~~~
*#!/bin/bash*
raxmlHPC-PTHREADS -f e -t tree.nwk -m GTRGAMMA -s aln_dna.phy -n REF -p 1984 -T 8
~~~

On the other hand, we used RAxML version 8.2.12 to compute backbone tree (branch lengths update only) and ML model parameters required for EPA-ng:

~~~
*#!/bin/bash*
raxmlHPC-PTHREADS -f e -t tree.nwk -m GTRGAMMA -s aln_dna.fa -n REF8 -p 1984 -T 8
~~~

For all large alignments with 10, 000 or more sequences in RNASim-VS, RAxML version 8.2.12 fail to run due to estimated gamma rate being either too small or too large. In those cases, we ran the following command to use GTRCAT model instead:

~~~
*#!/bin/bash*
raxmlHPC-PTHREADS -f e -t tree.nwk -m GTRCAT -s aln_dna.fa -n REF8 -p 1984 -T 8
~~~

For the same dataset, we used FastTree again for re-estimating Minimum Evolution based branch lengths and used that tree for performing placements using APPLES. FastTree is run with the following command:

~~~
*#!/bin/bash*
FastTreeMP -nosupport -nt -nome -noml -log tree.log
-intree tree.nwk < aln_dna.fa > tree_me.nwk
~~~

#### Performing placement

We performed phylogenetic placement of a query using pplacer version 1.1.alpha19-0-g807f6f3 with the following command:

~~~
*#!/bin/bash*
pplacer -m GTR -s RAxML_info.REF -t backbone.nwk -o query.jplace aln_dna.fa -j 1
~~~

We performed EPA-ng placements using version 0.3.5 with the following command:

~~~
*#!/bin/bash*
epa-ng --ref-msa aln_dna.fa --tree backbone.nwk --query query.fa --outdir.
--model RAxML_info.REF8 --redo -T 1
~~~

Except in our RNASim-VS, RNASim-QS and estimated alignments analyses, we used APPLES version 1.1.0 (can be found at https://github.com/balabanmetin/apples/releases) for placement. When alignment is not present, we performed the placement running the following command:

~~~
*#!/bin/bash*
python ∼/apples/least_squares_placement.py -t backbone.tree -d dist.mat -a FM
-s MLSE -p > apples.nwk
~~~

When alignment is present, we used the following command instead:

~~~
*#!/bin/bash*
python ∼/apples/least_squares_placement.py -t backbone.nwk -a aln_dna.phy
-a FM -s MLSE -p > apples.nwk
~~~

For RNASim-VS, RNASim-QS and estimated alignments analyses, we used APPLES version 1.2.0 and ran with the following command:

~~~
*#!/bin/bash*
python ∼/apples/run_apples.py -t backbone.nwk -s aln_dna.fa -q query.fa
-T $numcores -o apples.jplace
~~~

where $*numcores* depends on the number of cores designated for the analysis.

#### Working with estimated backbone and query-to backbone alignments

We created a version of SEPP within which APPLES integrated. This version is available at https://github.com/balabanmetin/sepp. We performed placement on RNAsim-AE dataset with 10,000 sequences using SEPP+APPLES with the following commmand:

~~~
*#!/bin/bash*
python ∼/sepp/run_sepp.py -t estimated_backbone.nwk -a estimated_aln.fa
-r RAxML_info.REF -f query.fa -pl apples -x 28 -A 1000 -o apples
~~~

On the same dataset, we ran SEPP in default mode using the following commmand:

~~~
*#!/bin/bash*
python ∼/sepp/run_sepp.py -t estimated_backbone.nwk -a estimated_aln.fa
-r RAxML_info.REF8 -f query.fa -pl pplacer -x 28 -A 1000 -P 1000 -o pplacer
~~~

